# Micro-CT Based Description of *Dorsetichthys bechei* (Actinopterygii: Teleostei): Cranial Anatomy of an Iconic Early Teleost

**DOI:** 10.1101/2024.05.10.593503

**Authors:** Jake Atterby, Matt Friedman, Sam Giles

## Abstract

Teleost fishes comprise more than 36,000 living vertebrate species and occupy a wide range of aquatic ecosystems. Despite their overwhelming success, extinct taxa outside of the living radiation—stem teleosts—remain poorly understood. *Dorsetichthys bechei* is an Early Jurassic stem teleost of historic importance. Although *D. bechei* is well described, its proximity to the teleost crown and relationships with other stem teleosts is challenging due to previous associations with the wastebasket assemblage known as “pholidophorids”. Here we present a new, three-dimensionally preserved specimen of *Dorsetichthys bechei* from the Blue Lias Formation (Sinemurian) Lyme Regis, UK. High-resolution CT scanning of this specimen plus additional material reveals previously unknown internal morphological information, including of the jaw and palate, hyoid arch, and gill skeleton. Importantly, this previously undescribed articulated specimen permits description of the external dermal skeleton and internal endoskeleton for the same individual, clarifying past accounts drawing on composite descriptions of two- and three-dimensionally preserved specimens. We confirm the presence of important teleost synapomorphies such as a mobile premaxilla and quadratojugal process. However, contrasting previous accounts, we find no evidence for dorsal and ventral hypohyals or a postarticular process of the lower jaw. We also confirm the presence of a prearticular. The presence of a mobile upper jaw, elaborate gill rakers and reduced dentition also support a possible facultative filter feeding ecology. These new anatomical data support removal of *D. bechei* from Pholidophoridae and Pholidophoriformes as currently defined but indicate that finer-scale patterns of relationships among more crownward members of the teleost stem are some way from being resolved.

## Introduction

Teleost fishes account for more than half of all extant vertebrate species, but the anatomy and relationships of taxa outside of the living radiation are poorly constrained despite an abundance of well-preserved material. Although past work identifies a broad range of Mesozoic actinopterygians as branching from the teleost stem (Patterson, 1973, 1975; Gardiner *et al*., 1996; Hurley *et al.,* 2007; Nursall, 2010; Arratia, 1997, 1999, 2013, 2017; Poyato-Ariza, 2015; Latimer & Giles, 2018), the anatomy and relationships of some key taxa are incompletely known (Arratia & Tintori, 1999; Arratia, 2001; Poyato-Ariza, 2015; Latimer & Giles, 2018), despite what can be exquisite preservation (e.g., Frey & Tischlinger, 2012; Cawley et al., 2019) and centuries of anatomical and taxonomic study (Agassiz, 1833 – 1843; Woodward, 1985).

“Pholidophoridae” (*sensu* Woodward, 1880), first established for the genus *Pholidophorus*, is an assemblage of modestly sized stem teleosts with a complicated taxonomic history. The importance of this group to understanding teleost evolution has been recognised for well over a century, with Woodward (1895) classifying pholidophorids as the earliest members of Isospondyli (early fossil teleosts and extant relatives). They later came to be classed within “holosteans.” A more inclusive group than that currently recognized (Grande, 2012), the “holosteans” of these authors also encompassed many other Mesozoic actinopterygians (e.g., aspidorhynchids, semionotids, pycnodonts) and the lineage leading to *Amia* (Romer, 1945). Although support for a direct link between pholidophorids and extant teleosts mounted (Woodward 1942, Rayner 1948, Saint-Seine, 1949; Gardiner, 1960; Griffith & Patterson, 1963; Nybelin, 1966; Patterson, 1968; Patterson, 1975), the large number of plesiomorphic characters displayed by the assemblage meant that influential figures such as Patterson initially resisted recognising them as teleosts rather than “holosteans” (Patterson 1967, 1968, 1973).

Early diagnoses of *Pholidophorus* largely cite vague and generic neopterygian characters (e.g., Agassiz, 1833-1843; Woodward, 1889; Lehman, 1966; Nybelin, 1966; Zambelli, 1986), leading to a taxonomic quagmire. At one point, the type genus *Pholidophorus* (Agassiz, 1832; 1837) contained over 40 species, most based on poorly preserved or inadequately described material (Woodward, 1895). Due in part to their position relative to more crownward parts of the tree, “pholidophorids” became a wastebasket taxon for early teleosts that retained a suite of primitive features such as rhombic ganoid scales (e.g., Patterson, 1975, pg. 281; Arratia, 2013, pg. 2-4). Perhaps understandably, “*Pholidophorus” bechei*, one of the better preserved “pholidophorids”, was erroneously considered the type species by Woodward and later authors (e.g. Rayner 1948, pg. 318; Nybelin 1966, pg. 32; see discussion in Arratia 2000), and both its dermal skeleton and neurocranium were described in detail. Several efforts have been made to revise the taxonomy of Pholidophoridae, with many species being reassigned to new genera (e.g., Deecke, 1889; Woodward, 1941; Nybelin 1966; Gaudant, 1978; Arratia, 2000). Work to tackle this problem culminated in Arratia’s (2013) comprehensive anatomical, taxonomic, and phylogenetic reassessment of pholidophorids and associated taxa, which resolved a monophyletic remnant of pholidophorids as sister to *Eurycormus* and all remaining teleosts (Arratia, 2013). Today, Pholidophoridae (*sensu* Arratia, 2013) is restricted to around 15 genera spanning the Late Triassic (Ladinian) of Europe and China.

Significantly, Arratia’s revision excluded “*Pholidophorus*” *bechei* from Pholidophoridae, removing it to its own monotypic genus *Dorsetichthys*, within a monotypic order Dorsetichthyiformes and family Dorsetichthyidae (Arratia, 2013; 2017). *Dorsetichthys* was recovered as closer to the teleost crown than pholidophorids, a placement supported by seven synapomorphies: a skull roof with an orbital region slightly narrower than the postorbital region, small teeth on the parasphenoid, a well-developed postarticular process of the lower jaw, absence of a prearticular, absence of serrated appendages, a pectoral propterygium fused with the first lepidotrich, and three rows of rhomboidal mid-flank ganoid scales between the postcleithra and mid-length of the body (Arratia, 2013). The removal of *Dorsetichthys* from pholidophorids also has implications for interpreting broader patterns of relationships on the teleost stem: this taxon has been functionally emblematic of pholidophorids for more than a century (Arratia, 1999). Because it has been described in the most detail, it is often the only “pholidophorid” taxon included in analyses of teleost or actinopterygian relationships (e.g., Patterson and Rosen, 1977; Olsen, 1984; Grande and Bemis, 1998; Arratia, 1997, 1999, 2001; Hurley et al., 2007; Xu & Wu, 2012; Poyato-Ariza, 2015; Thies & Waschkewitz, 2015; Gibson, 2016). Consequently, it is unclear to what extent the pholidophorid terminal in some past analyses represents a composite taxon, and to what extent valid pholidophorid taxa are represented.

While the cranial dermal (Nybelin, 1966) and endoskeletal (Patterson, 1975) anatomy of *Dorsetichthys* has been studied in great detail in comparison to most other stem teleosts, these accounts are typically taken from different individuals, with less clear aspects of anatomy supplemented by comparison to taxa such as *Pholidophorus latiusculus* or *Leptolepis* (e.g. Rayner, 1948). Rayner (1948) described the dermal skeleton, braincase, hyomandibula and palate and Patterson (1975) the neurocranium from isolated material, the species determination of one braincase (NMHUK PV P 51682) being noted by Patterson (1975: pg. 283) as uncertain due to the lack of other pholidophorids in which the braincase was known. Likewise, accounts of the dermal skeleton made by Nybelin (1966) are based on two-dimensional, laterally compressed specimens in which the endoskeleton is largely unknown, and the internal aspect of many dermal bones cannot be assessed. Most recently, Giles *et al*. (2018) used CT scanning to describe the endocast and bony labyrinth of another isolated braincase, NMHUK PV P 51682 (referred to incorrectly by Giles *et al*. as NHMUK PV P 1052), supplementing past preliminary descriptions of the endocranial cavity that were based on sagittal sections (e.g., Patterson, 1975). While our understanding of the anatomy of this taxon is comprehensive, it assumes that these isolated elements scattered across the dermal and endoskeleton belong to the same taxon.

Here, we present a complete, articulated, previously undescribed individual of *Dorsetichthys bechei* and a redescription of a specimen previously studied by Rayner (1948) and Patterson (1975). Through high resolution CT scanning, we describe both the dermal and endoskeletal anatomy of these specimens. In doing so, we clarify ambiguity regarding the compositing of isolated material and providing new details on its internal morphology. We then consider the consequences of this refined anatomical picture of *Dorsetichthys* for its placement relative to pholidophorids and other stem teleosts.

## Materials & Methods

### Institutional Abbreviations

NHMUK = Natural History Museum, London, UK.

### Materials

NHMUK PV P 73995 is a heavily pyritised, three-dimensionally preserved specimen, the history of which is poorly known. It was recently located among the cabinets containing specimens from the NHMUK collections as well as borrowed fossils studied by Colin Patterson; this was effectively his “working collection” of material. The specimen was kept with a hand-written note dated 1888 and signed by William Davies, but the stratigraphic origin and provenance of the specimen is unclear. Despite no indication on Davies’ associated note, the matrix and mineralisation are consistent with fossils from the Blue Lias Formation, Lyme Regis, UK (Lord & Davis, 2010). Upon its rediscovery, the specimen was provisionally identified as *Dorsetichthys bechei* and was formally registered and assigned a catalogue number in 2016.

NHMUK PV P 1052 is a mechanically prepared braincase from the Early Jurassic (Sinemurian) Charmouth (UK), with some associated dermal elements. No precise geological provenance is given but it is assumed to be from the Charmouth Mudstone Formation. Unlike NHMUK P 73995, NHMUK PV P 1052 has a well-documented history of study and description and has informed several composite reconstructions by Rayner (1948) and Patterson (1975). It was originally identified as *Pholidophorus bechei*, and later removed to *Dorsetichthys bechei* (Arratia, 2013).

Several additional specimens of *D. bechei* were also examined for comparison of the braincase and dermal skeleton, and to investigate the possibility of intraspecific variation in the circumorbital series: NHMUK OR 19010, NHMUK OR 38107, NHMUK OR 38109, NHMUK OR 39859, NHMUK PV P 3589a, NHMUK PV P 1051, NHMUK PV P 1052a, NHMUK PV P 3586c, NHMUK OR 154, NHMUK OR 38535 and NHMUK OR 1885. Each specimen is attributed to the Lower Lias, most likely corresponding to the modern Charmouth Mudstone Formation, but their precise provenance is poorly recorded. NHMUK OR 19010 comprises a three-dimensionally preserved cranium and complete body, which also preserves some relief. The braincase and associated cranium can be separated from the rest of the specimen and was previously studied by Patterson (1975). The remaining specimens comprise laterally compressed dermal skeletons of varying completeness which have also previously informed composite descriptions by Rayner (1948) and Nybelin (1966).

### CT Scanning and Imaging

NHMUK PV P 73995 was scanned at the CTEES Facility, University of Michigan using a Nikon XTH 225 ST with a 0.85 mm Cu filter; current = 180 μA; voltage = 195 kV; exposure = 1000 ms and 4 frames per projection, resulting in a voxel size of 21.68 μm. NHMUK PV P 1052 was scanned at the Imaging and Analysis Centre, NHMUK using a Zeiss Versa 520 micro-CT scanner with a HE2 filter (unspecified thickness of glass); current = 63 μA; voltage = 160 kV; exposure = 8000 ms and 1 frame per projection, resulting in a voxel size of 20.59 μm.

Resulting image stacks were cropped in ImageJ (Schneider *et al*., 2012) and segmented in Mimics v.25 (biomedical.materialise.com/mimics; Materialise, Leuven, Belgium). Renders of 3D models were captured using Blender v. 3.3.0 (blender.org) and interpretive drawings were created in Adobe Illustrator (adobe.com/products/illustrator).

## Systematic Palaeontology

Osteichthyes Huxley, 1880

Actinopterygii Cope, 1887

Neopterygii Regan, 1923

Teleostei Müller, 1844

Dorsetichthyiformes Nelson, Grande & Arratia, 2016

Dorsetichthyidae Nelson, Grande & Arratia, 2016

*Dorsetichthys* Arratia, 2013

*Dorsetichthys bechei* (Agassiz, 1837)

### Holotype

PAL-J.003065, Oxford University Museum of Natural History.

## Diagnosis (emended from Arratia, 2013)

Neopterygian diagnosed by the following unique combination of characters: partially fused skull roof plate; five infraorbital bones between the antorbital and dermosphenotic; one or two suborbital bones; two or three supraorbital bones; thin lateral strut extending across the sub-temporal fossa of the braincase; mobile premaxilla and maxilla articulating with the ethmoid; four ural neural arches modified as uroneurals; hypurals divided into dorsal and ventral groups.

## Description: NHMUK PV P 73995

### General morphology

NHMUK PV P 73995 (Fig. 1) consists of a near-complete skull, as well as a partial postcranium with scales and a pectoral fin. The left side of the skull and the skull roof have been exposed following mechanical preparation, but several bones on the exposed side are displaced and/or damaged. The right side of the skull, however, is preserved within the matrix and almost every dermal bone is present and in articulation. The entire fossil has undergone secondary mineralisation by pyrite, giving most of the specimen a golden, metallic lustre and relatively high contrast in tomograms.

**Figure 1:**
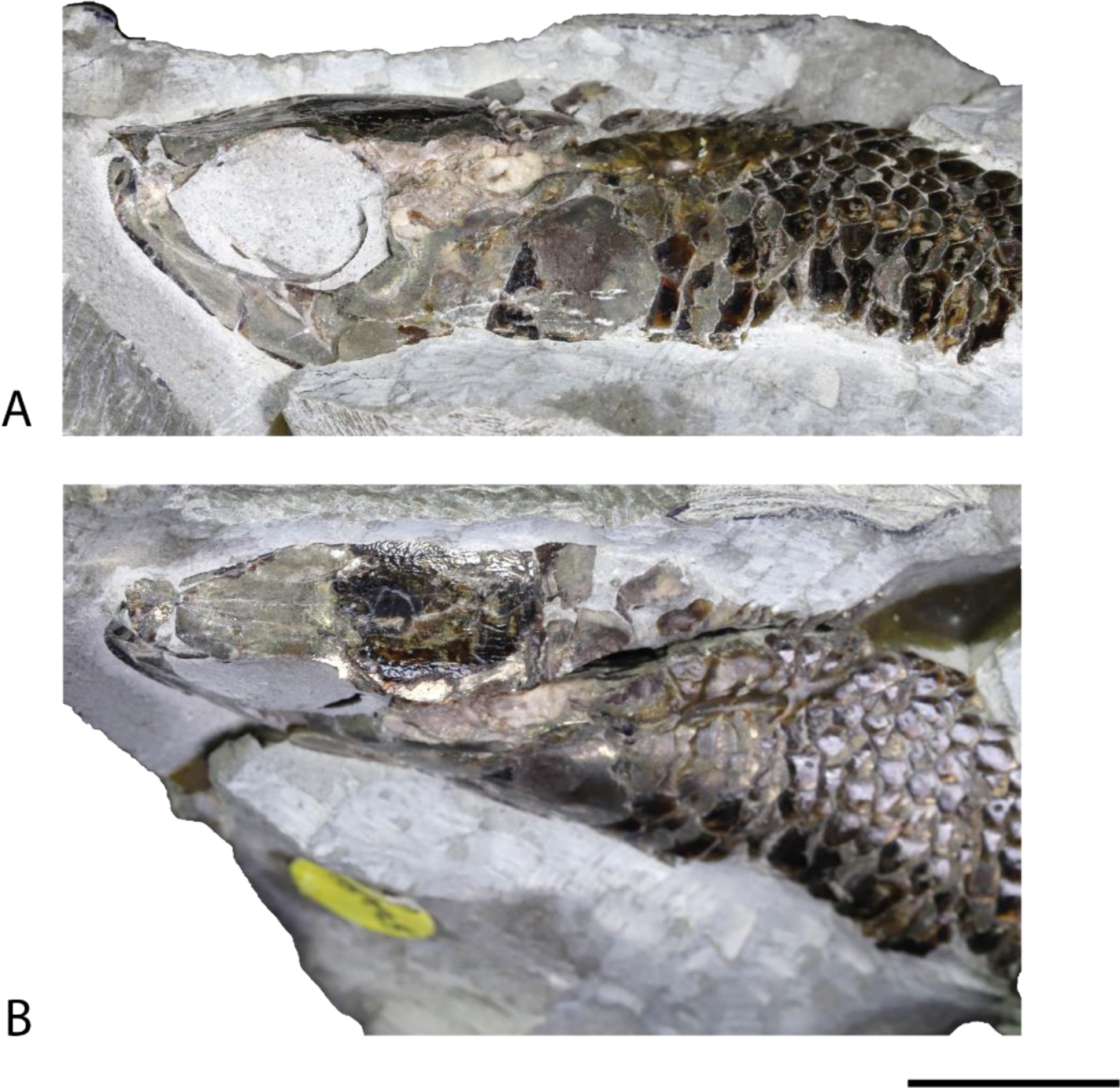
*Dorsetichthys bechei* NHMUK PV P 73995. Scale = 10 mm.

### Skull Roof

The dermal plates of the skull roof are partially preserved. Overall, the skull roof is broadest posteriorly, tapering abruptly above the postorbital margin and towards the snout. The broadest section of the skull roof is roughly four times wider than the anterior region. The skull roof comprises paired frontals, parietals and dermopterotics (fr, pa, dpt, Figs. 2B,D, 4B). Sutures between individual bones are difficult to trace in tomograms but visible on the specimen. They are largely straight, or gently curved with the exception of the strongly sinusoid suture between the midsection of the frontals (Fig. 2B). The broader, posterior portion of the skull roof is damaged on the left side, but the right is well preserved within the matrix, exhibiting anterior, middle, and posterior pit lines (pl, Fig. 2D). Enclosed supraorbital canals (so.c, Fig. 2D) extend along the full length of the plate to the nasal region and are exposed as moulds anterior to the preserved portion of bone. Each canal opens to the surface via irregularly spaced pores that accommodate simple tubules. The right infraorbital canal (io.c, Fig. 2D) is fully enclosed in a ridge on the ventral margin of the skull roof plate, extending along its lateral margin and exiting the posterior margin into the extrascapular (exc, Fig. 2B, D).

**Figure 2:**
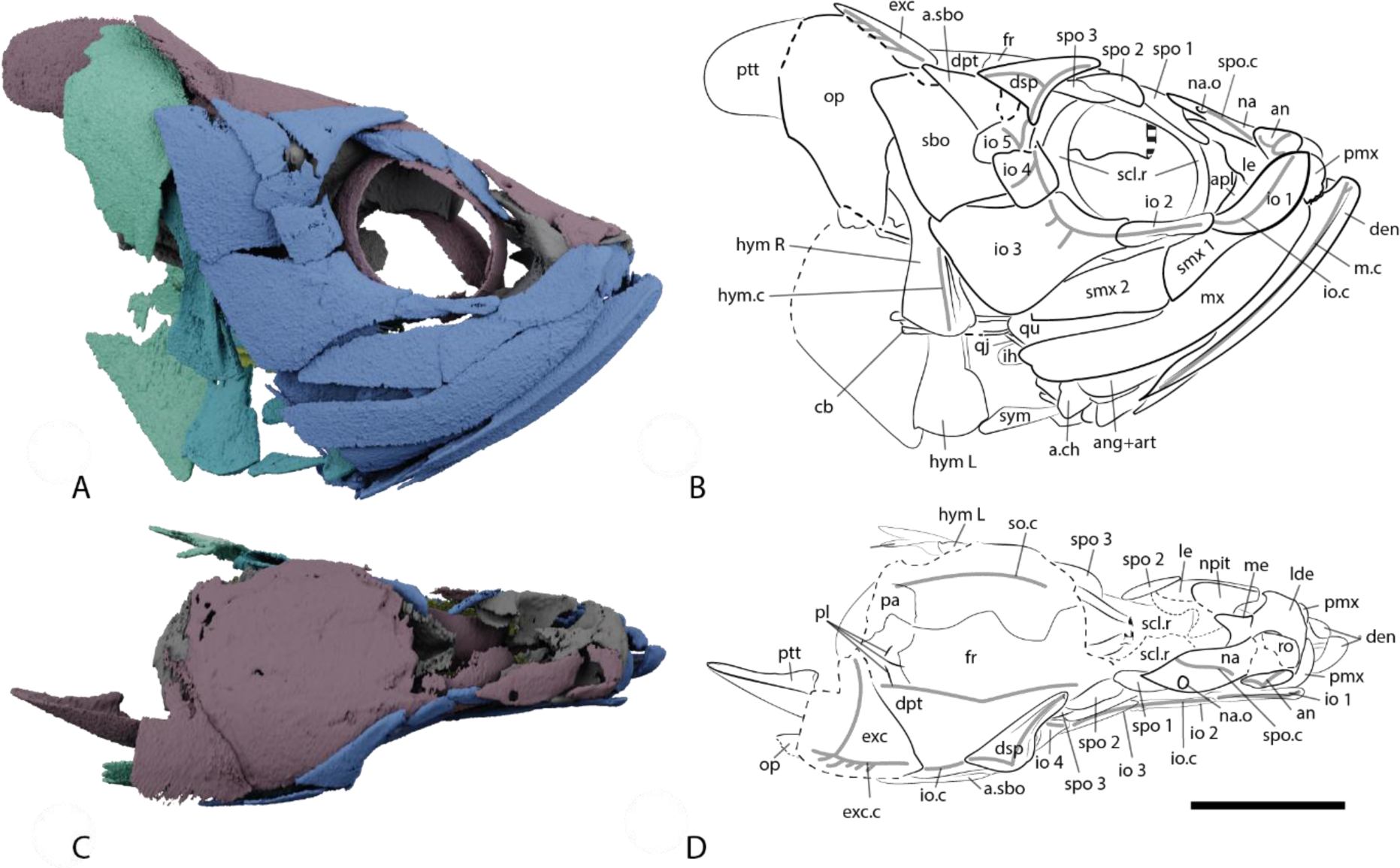
The skull of *Dorsetichthys bechei* NHMUK PV P 73995. (A) Render and (B) interpretive drawing in right lateral view. (C) Render and (D) interpretive drawing in dorsal view. Colour coding: blue, cheek and jaw; purple, skull roof, dermal shoulder girdle and sclerotic ossicle; grey, braincase; turquoise, hyomandibula; light green, operculogular system; yellow, gill skeleton. Sensory lines indicated in grey. Scale bar = 10 mm. *Abbreviations: an, antorbital; ang+art, fused angular and articular, apl, autopalatine; angular; a.sbo, accessory suborbital; cb, ceratobranchial; a.ch, anterior ceratohyal; den, dentary; dpt, dermopterotic; dsp, dermosphenotic; exc, extrascapular; exc.c, extrascapular canal; fr, frontal; hym, hyomandibula; ih, interhyal; io.c, infraorbital canal; io 1-5, infraorbital bones; le, lateral ethmoid; lde, laterodermethmoid; me, mesethmoid; mx, maxilla; m.c, mandibular canal; na, nasal; na.o, nasal opening; npit, nasal pit; op, opercular; pa, parietal; pl, pit lines; pmx, premaxilla; ptt, posttemporal; qu, quadrate; qj quadratojugal process; sbo, suborbital; scl.r, sclerotic ring; smx 1-2, supramaxillae; so.c, supraorbital canal; spo 1-3, supraorbital bones; spo.c, supraorbital canal; sym, symplectic*.

The right nasal (na, Figs. 2B, D, 6B) is well preserved, although the left nasal is missing entirely. It is sub-rectangular with a stepped, tapered posterior extension that lies parallel to the supraorbital series. It carries the supraorbital sensory canal (spo.c, Fig. 2B, D) along its entire length and bears the posterior nasal opening (na.o, Figs. 2B, D, 6B). Though not preserved in this specimen, previous descriptions (Nybelin, 1966; pg. 360) show the frontals separating the left and right nasals. The antorbital (an, Figs. 2B, D, 6B) is small and triangular, forming the most anterior portion of the orbital margin, partially overlapping the nasal and lying anterodorsally to the first infraorbital. Two or three pores from the infraorbital canal (io.c, Figs. 2B, D, 6B) open onto the lateral surface of the antorbital. The canal bifurcates within the bone, with a dorsal branch continuing into the nasal, and the ethmoid commissure continuing anteriorly into a small fragment of the right rostral bone (ro, Fig. 2D). Despite being poorly preserved, the rostral fragment appears to be *in situ*. The nasal, rostral and antorbital bones all partially overlap in the ethmoid region, and the right nasal bone has partly subsided into the underlying nasal pit.

### Braincase

The bulk of the braincase consists of a single ossification encompassing the occipital, otic, orbitotemporal and basisphenoid regions; separate ossifications or ossification centres cannot be identified. However, both the orbitosphenoid and ethmoid regions are independently ossified, and entirely separated from the other portions of the braincase that presumably would have been filled by cartilage in life.

#### Occipital Region

The occipital region is well preserved and relatively well ossified, with a complete lining of perichondral bone and moderate development of endochondral bone posteriorly. However, ossification is insufficient to trace or identify some canals for nerves and vessels. The occipital region is comparatively narrow, and bounded anteriorly by a shallow groove on the lateral margin of the braincase that marks the closed otoccipital fissure (fotc, Fig. 3G). In posterior view (Fig. 3H), the region corresponding to the supraoccipital forms a pronounced, hourglass-shaped, vertical ridge of bone flanked by shallow depressions. Contra Patterson (1975, pg. 307), there is no evidence that the medial portion of the “supraoccipital region” has a dermal component as it passes beneath the parietals. An oval opening for the foramen magnum (fm, Fig. 3H) lies immediately ventral to the supraoccipital ridge. Its ventral margin is thickened into overhanging posteroventral processes, homologous with the occipital condyle (occ, Fig. 3H, K), and likely articulated with the intercalaries of the first vertebra (Patterson, 1975). The inner walls of the foramen magnum narrow anteriorly and appear to lack any distinct foramina. More ventrally, the rounded notochordal opening (not, Fig. 3H), extends into the braincase as a conical canal that constricts rapidly and terminates anteriorly at the level of the saccular recess. A C-shaped aortic notch forms the most ventral feature on the posterior face of the occipital region, framing the walls of a deep, open groove for the dorsal aorta (aort, Fig. 3H, K, L) that extends along the ventral surface of the basiocciput. In agreement with Patterson (1975), we find no evidence for an enclosed aortic canal. The posterior margin of the aortic notch extends posteriorly to the level of the occipital condyle, forming the basal point of articulation with the vertebrae.

**Figure 3:**
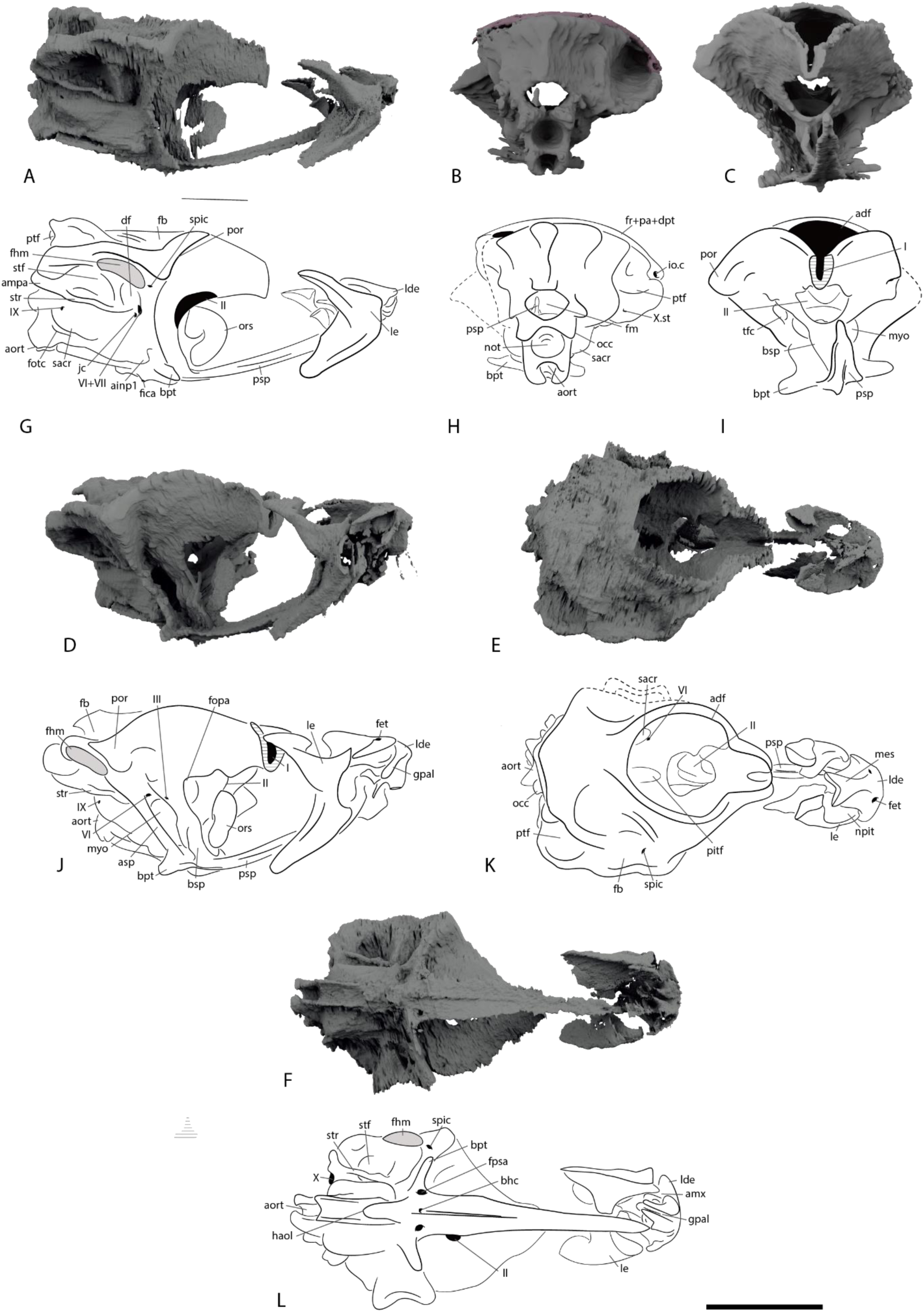
The braincase of *Dorsetichthys bechei* NHMUK PV P 73995. Renders in (A) right lateral, (B) posterior, (C) anterior, (D) anterolateral, (E) dorsal, and (F) ventral view. Interpretive drawings in (G) right lateral, (H) posterior, (I) anterior, (J) anterolateral, (K) dorsal, and (L) ventral view. Scale bar = 10 mm. Abbreviations*: adf, anterior dorsal fontanelle; aort, aortic notch; bhc, buccohypophysial canal; bpt, basipterygoid process; bsp, basisphenoid; df, dilatator fossa; epsa, efferent pseudobranchial artery; fb, fossa bridgei; fhm, hyomandibular facet; fm, foramen magnum; fotc, otoccipital fissure; fr+pa+dpt, fused frontal, parietal and dermopterotic (skull roof); haol, housing of aortic ligament; ica, internal carotid artery; myo, posterior myodome; not, notochord; pf, pituitary fossa; por, post-orbital process; psp, parasphenoid; ptf, post-temporal fossa; sacr, saccular recess; socc, supraoccipital; spic, spiracular canal; str, lateral strut; tfr, trigemino-facial recess; II, optic fenestra; VI, foramen of abducens nerve; VII, foramen of facial nerve X, foramen of vagus nerve*.

Anterior to the closed otoccipital fissure, a ventrally directed elliptical opening marks the exit of the vagus nerve (X, Fig. 3L). The smaller supratemporal branch of the vagus nerve appears to exit anterolaterally (X.st, Fig. 3H), although the canal itself is hard to trace internally. A swelling dorsal to the exit of the vagus nerve, in the region formed by the exoccipital, corresponds to the position of the posterior ampulla of the inner ear (ampa, Fig. 3G). Foramina for the occipital nerve and posterior cerebral vein cannot be identified. The posterior myodome (myo, Fig. 3I, J), which originates in the posterior orbital wall, extends well into the occipital region, narrowing posteriorly and terminating level with the posterior margin of the saccular cavities. There is no posterior dorsal fontanelle or vestibular fontanelle, and the ventral otic fissure is also closed.

#### Otic Region

The otic region represents the widest portion of the braincase, approximately 1/3 wider than the occipital region. Much of this region is poorly ossified, with only thin perichondral bone forming most margins, and no endochondral development. As with the occipital region, this makes the path of canals difficult to follow. The endocranial roof is almost entirely unossified and is instead open as a large anterior dorsal fontanelle (adf, Fig, 3I, K), which extends to the anterior margin of the orbital region. Several fractures disrupt the left side of the otic region, but the right side is well preserved, revealing several fossae. The post-temporal fossa (ptf, Fig. 3G, H, K) is wide, rounded, and gradually narrows anteriorly towards the complex fossa bridgei (fb, Fig. 3G, J, K). The fossa bridgei is very broad, with a shallow, convex dorsal margin, narrowly separated from the anterior dorsal fontanelle and cranial cavity by the housing of the infraorbital canals in the skull roof. The fossa bridgei does not appear to be confluent with the post-temporal fossa, although ossification between them is very thin. Both depressions lack an endocranial roof and are overlain exclusively by the dermal skull roof. At its deepest point, the floor of the fossa bridgei is connected to the dorsal opening of the spiracular canal (spic, Fig. 3G, K). The spiracular canal is vertically oriented and enclosed in bone within the postorbital process (por, Fig. 3G, I, J), re-emerging ventrally into a groove roughly level with the anterior foramen of the abducens nerve. No bifurcations or branching rami are resolved as extending from the spiracular canal, consistent with Patterson (1975).

The hyoid facet (fhm, Fig. 3G, J, L) and sub-temporal fossa (stf, Fig. 3G, L) lie ventral to the post-temporal fossa and fossa bridgei. The hyoid facet is fairly shallow, oriented sub-horizontally, and lined with perichondral bone. Anteroventrally, the sub-temporal fossa forms a considerably deeper recess, partially delineated from the hyomandibular facet and post-temporal fossa by a thick ridge of bone that houses the horizontal semicircular canal. A thin lateral strut (str, Fig. 3G, J, L) of endochondral bone extends from the intercalar region across the full length of sub-temporal fossa and reconnects with the braincase on the underside of the utricular recess.

The jugular groove passes ventral to and parallel with the lateral strut. It is shallow without sharply defined margins but appears to extend posteriorly from the glossopharyngeal foramen (IX, Fig. 3 J, G) and across the external face of the saccular chamber (sacr, Fig. 3G, H) before piercing the postorbital process and entering the jugular canal (jc, Fig. 3G). The foramen for the facial nerve (VII, Fig. 3G) opens laterally into the floor of the jugular groove. The anterior wall of the shallow dilatator fossa (df, Fig. 3G) contributes to part of the anterior wall of the otic region. A smooth and rounded swelling, which narrows anteriorly, corresponds the saccular chamber and forms much of the ventral portion of the lateral face of the otic region. A stout, sub-triangular anterolateral process (ainp1, Fig. 3G) anterior to the saccular recess represents the articulation point for the infrapharyngobranchial of the first gill arch.

#### Orbitotemporal Region

The orbitotemporal region is relatively narrow and poorly ossified anteriorly. Posteriorly, the orbitotemporal region is separated from the otic region by a steep wall formed by the postorbital process, lateral commissure, and ascending process of the parasphenoid. The posterior orbital wall is well ossified and opens ventrally to the facial recess and median posterior myodome. Several shallow fossae occupy the posterior corner of the orbital cavity. The largest, the trigemino-facial recess (tfc, Fig. 3I), forms a short longitudinal tunnel in the wall of the postorbital process. The chamber includes the dorsal openings of the abducens nerve and the palatine branch of the facial nerve (VII, Fig. 3G), which exit the cranial cavity to pierce the concave roof of the large, median posterior myodome (myo, Fig. 3I, J, K). Though it forms a single median chamber, its anterior opening is bisected medially by the vertical pillar of the basisphenoid (bsp, Fig. 3J, K). The basisphenoid is notably reduced but frames the large median optic foramen (II, Fig. 3J, K), as well as the subtriangular pituitary fossa (pitf, Fig. 3k) which allows the posterior myodome to communicate with the optic foramen above it.

The median optic foramen (II, Fig. 3G, I, J, K, L) is large. A small notch on its dorsal margin would likely have accommodated the optic artery (fopa, Fig. 3J). The foramen for the trochlear nerve is hard to identify but may be present as a perforation halfway up the left orbital wall (IV, Fig. 3K). A small opening adjacent to the optic foramen may represent the oculomotor foramen (III, Fig. 3J). There is a short gap between the postorbital ossification and the lateral ethmoids.

#### Ethmoid

The ethmoid region is mostly complete and articulated, although some ossifications have shifted relative to each other. Overall, the ethmoid contributes to a rounded snout, with large paired nasal pits (npit, Figs. 3K, 4B, D, F). The ethmoid complex includes partially fused paired lateral dermethmoids (lde, Figs. 3J, K, L, 4B, D, F), lateral ethmoids (le, Fig. 4) and a median mesethmoid (mes, Figs. 3K, 4B, F). Sutures between these ossifications are difficult to identify, even in tomograms. The mesethmoid has a broad, subtriangular dorsal surface, which tapers ventrally to a thick nasal septum (nse, Fig. 4B, D) that forms the medial walls of the nasal pits. The ventral surface of the mesethmoid bears a very deep but narrow median groove (gpal, Figs. 3L, 4H, J). This incision likely transmitted the palatine nerve to the relatively shallower palatine notch (npal, Fig. 4B) between the articular surfaces for the maxillae and premaxillae (see below). Although the ethmoid is slightly offset from the remaining braincase, the anterior margin of the parasphenoid or vomer (the latter of which is not preserved) also likely extended into the posterior margin of the groove. The groove is flanked by lateral fossae, which may have accommodated the vomer.

**Figure 4:**
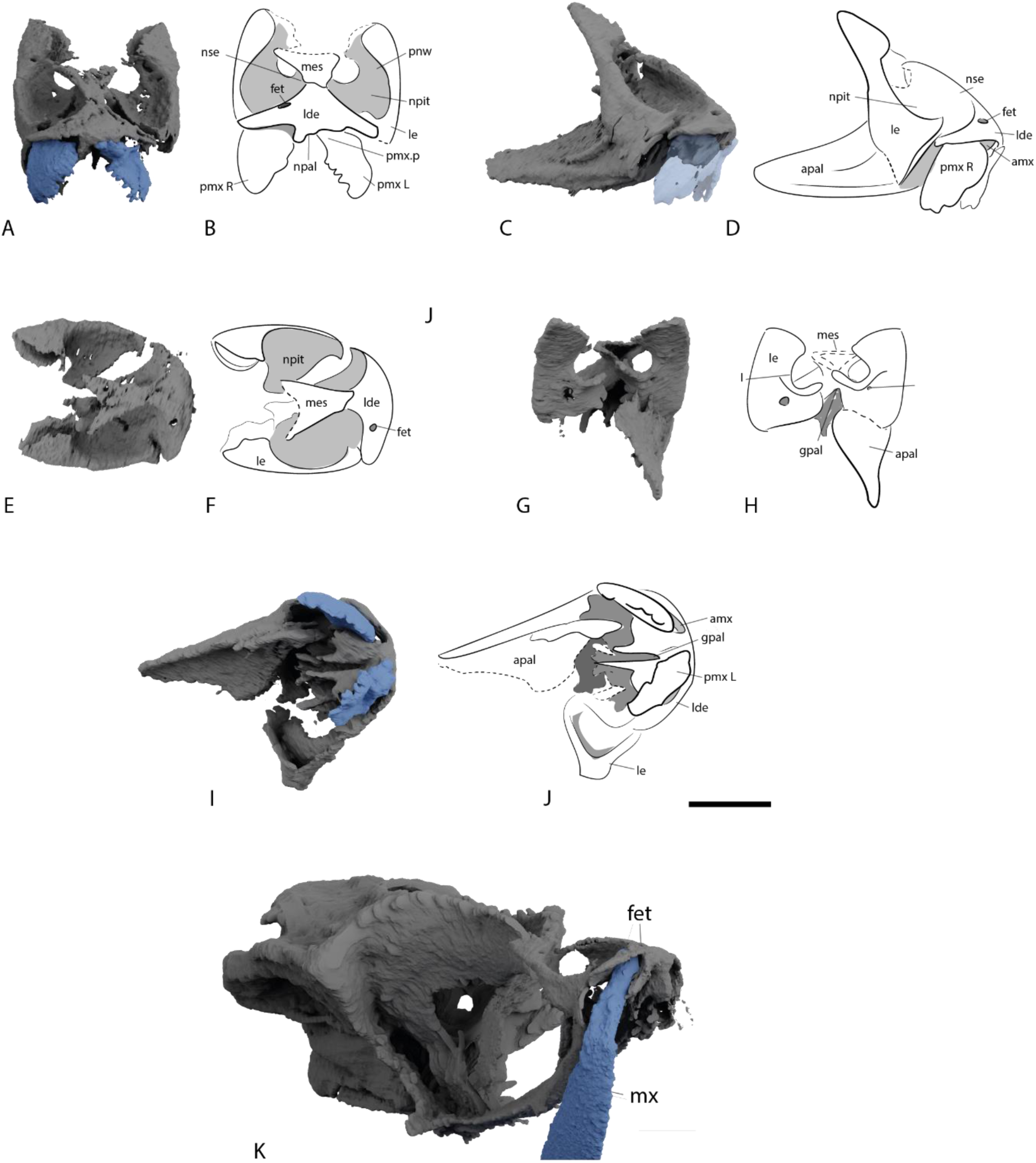
The ethmoid of *Dorsetichthys bechei* NHMUK PV P 73995. Renders in (A) anterior, (C) right lateral, (E) dorsal, (G) posterior, (I) ventral view, and interpretive drawings in (B) anterior, (D) right lateral, (F) dorsal, (H) posterior, (J) ventral view. Render (K) of right maxilla articulating with the ethmoid via the fenestra. Colour coding: blue, jaws; grey, braincase. Scale bar = 5 mm. Abbreviations: *amx, articular surface for maxilla and premaxilla; apal, autopalatine; fet. ethmoid fenestrae for articulation with maxilla, gpal, groove for palatine nerve; lde, laterodermethmoid; le, lateral ethmoid; mes, mesethmoid; mx, maxilla; npal, notch for palatine nerve; npit, nasal pit; nse, nasal septum; pmx, premaxilla; pmx.p, premaxillary process*.

The lateral dermethmoids are fused along the midline and extend posteriorly to contribute to the floor of the nasal pit and anteriorly beneath the rostral (Fig. 2D) to contribute to the snout. Teeth appear to be absent from this anterior margin. A large, rounded foramen on the anterodorsal face of the lateral dermethmoid (fet, Figs. 3J, K, 4B, F) resembles that previously observed in “*Pholidophorus” germanicus*, where it was described as “an unknown foramen or pit” by Patterson (1975: pg. 477). This opening communicates ventrally with a large socket (amx, Figs. 3L, 4J), bounded laterally and dorsally by the lateral dermethmoid and medially by the mesethmoid. The more dorsal part of this socket accommodates the anterior process of the maxilla—which extends directly to the foramen on the face of the lateral dermethmoid—and the more ventral the anterior process of the premaxilla, enabling both bones to articulate with the braincase and the upper jaw to protrude. It is likely that this feature is widely distrusted amongst other taxa branching from similar parts of the teleost stem, such as *Ichthyokentema* (Griffith & Patterson, 1963) and *Leptolepis* (Patterson, 1975). However, dermal bones obscure the position of a potential facet in these examples.

Posteriorly, the lateral ethmoids form the remainder of the floor and walls of the nasal pit, as well as the postnasal wall (and thus the anterior face of the orbits). Multiple small foramina perforate the postnasal wall, and likely correspond to orbitonasal vessels. A much larger, rounded opening, the roof of which is unossified, transmitted the olfactory nerve from the orbit into the nasal capsules (I, Figs. 3I, J, 4D). This opening may also have been confluent with the anterior myodome.

#### Parasphenoid

The parasphenoid (psp, Fig. 3G, I, J, K, L) spans almost the entire length of the braincase, terminating posteriorly just before the occiput and anteriorly beneath the ethmoid. A deep, median groove for the dorsal aorta occupies the ventral surface of the parasphenoid anterior to the aortic notch (aort, Fig. 3I). The groove is widest posteriorly, and tapers gradually anteriorly towards a narrow, deep canal for accommodating the aortic ligament (haol, Fig. 3J, K, L). Anterior to this opening, the parasphenoid broadens slightly and the dorsal aorta bifurcates; paired, shallow grooves on the ventrolateral face of the parasphenoid carry the lateral dorsal aortae forward. Further anteriorly, the parasphenoid bears a ventral keel, along with ascending processes that extend dorsally to meet the postorbital process. Slender, elongate dermal basipterygoid processes (bpt, Fig. 3G, I, J, L) project anterolaterally from the main body of the parasphenoid at a roughly 45° angle. The internal carotids pierce their posterior margin (fica, Fig. 3G), while the efferent pseudobranchial arteries (fpsa, Fig. 3L) occupy large, rounded foramina near their anterior margin.

The ventrally-directed opening of the buccohypophysial canal (bhc, Fig. 3L) is positioned directly between the efferent pseudobranchial foramina; it is poorly preserved in this specimen, but its presence is corroborated by NHMUK PV P 1052. The buccohypophysial canal is surrounded by a raised, rounded rim of well-ossified bone on the ventral surface of the parasphenoid. This forms the posterior margin of a ventral keel that runs along the midline of the anterior corpus, fading out anteriorly. The anterior corpus of the parasphenoid is narrow and elongate, triangular in profile, and extends well anterior to the orbits. Although too poorly resolved to segment, examination of the tomograms suggests the parasphenoid continues ventral to the ethmoid, inserting into the deep groove on the underside of the mesethmoid (gpal, Fig. 3J, L, 4H, J). No dentition is visible on the ventral surface, but minute teeth have previously been described in the taxon (Patterson, 1975, pg. 328, 520), and are likely beyond the resolution of the scan.

### Endocast

Despite poor mineralisation of the walls of the cranial cavity, the medial and right side of the endocast can be reconstructed (Fig. 5). The left side is incomplete due to fractures. Overall, the endocast is short, rounded, and robust. Much of its dorsal surface, from the anterior limit to the level of the crus commune, is open to the anterior dorsal fontanelle.

**Figure 5:**
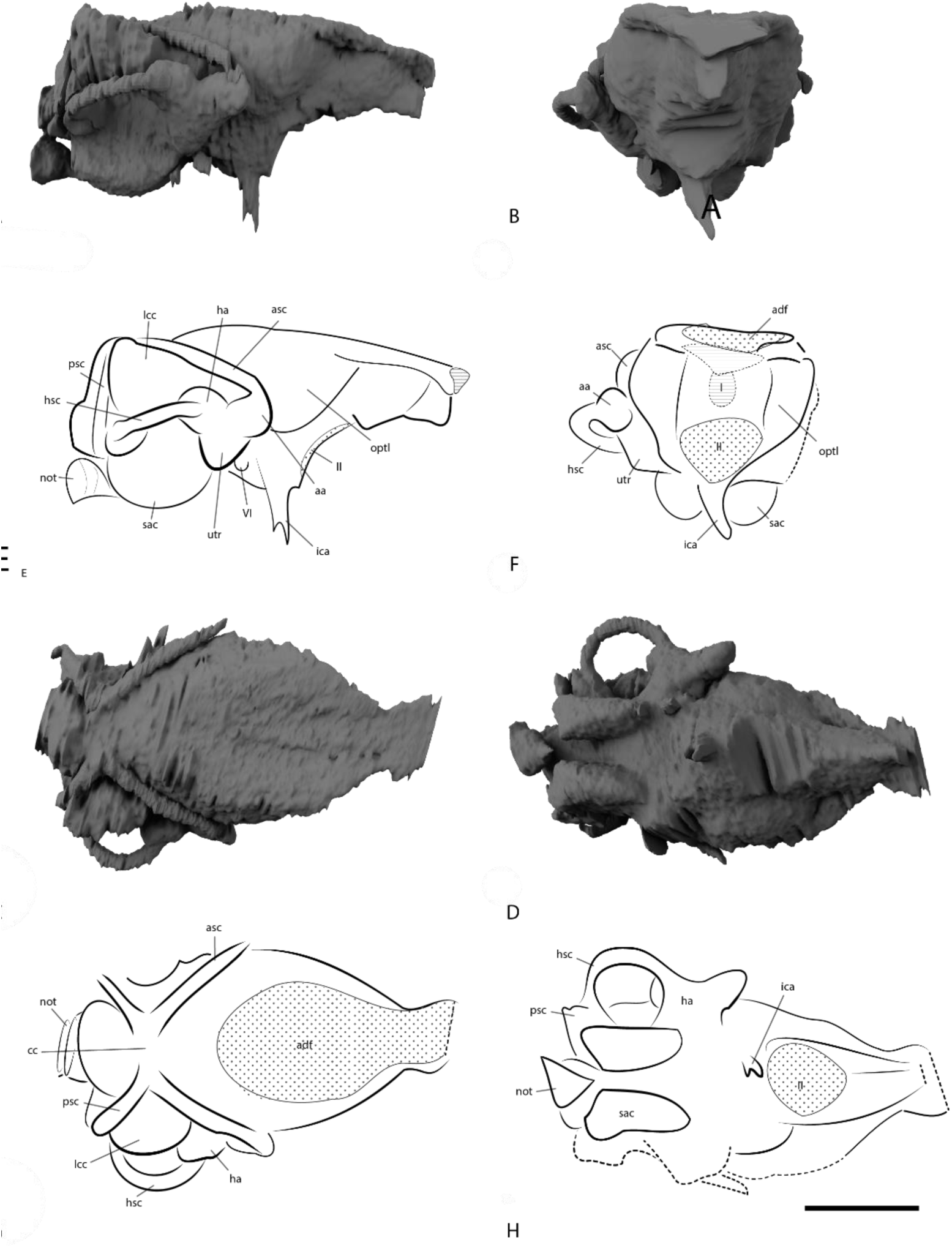
The endocast of *Dorsetichthys bechei* NHMUK PV P 73995. Renders in (A) right lateral, (B) anterior, (C) dorsal and (D) ventral view. Interpretive drawings in (E) right lateral, (F) anterior, (G) dorsal, and (H) ventral view. Scale bar = 5 mm. Abbreviations*: aa, anterior ampulla; asc, anterior semicircular canal; aur, cerebellar auricles cc, crus commune; ha, horizontal ampulla; hsc, horizontal semicircular canal; ica, internal carotid artery; lcc, lateral cranial canal; not, notochordal canal; optl, optic lobe; psc, posterior semicircular canal; sac, saccular; utr, utricular; II, optic nerve; VI, abducens nerve; X, vagus nerve*.

The hindbrain region occupies around a third of the length of the endocast and has a mineralised roof. It is shallower than the rest of the endocast and restricted ventral to the anterior semicircular canal. The cerebellar auricles are very small, and difficult to distinguish from the rest of the endocavity. The root of the vagus nerve is oval, and exits the endocranial cavity ventrally. It is initially confluent with the posterior ampulla, but becomes separated by a very thin sheet of bone more distally from the endocranial cavity. There is no distinct cerebellar corpus.

Almost the entire midbrain region of the endocast is occupied by the optic tectum. The paired swellings for the optic lobes (optl, Fig. 5E, F) are deep and rounded, and wider than both the cerebellar and telencephalic regions of the endocast. However, the dorsal margin of the optic lobes is unclear on account of the open anterior dorsal fontanelle. The abducens nerve (VI, Fig. 5E) leaves the cranial cavity in this region, piercing the postorbital process before opening ventrally into the roof of the myodome.

Very little of the region corresponding to the forebrain can be described, largely due to poor mineralisation of the orbital walls of the endocavity. The area for the forebrain is the narrowest portion of the endocast, with convex lateral walls. Its anteroventral margin is continuous with the large, median optic foramen (II, Fig. 5E, F, H). Ventral to the optic nerve is a narrow canal for the internal carotid artery (ica, Fig. 5E, F, H), which enters the basisphenoid via the pituitary fossa (pitf, Fig. 3L). The proportions of the skull suggest elongated olfactory tracts, but no other features can be inferred.

All three semicircular canals are preserved and fully enclosed in bone. The canals are all equally narrow gauge, but of varying lengths. The anterior semicircular canal (asc, fig. 5E, G) is long and mostly straight, and closely applied to the cerebellar region of the endocranial cavity. The horizontal semicircular canal (hsc, Fig. 5E, G, H) is tightly curved, forming a half circle that enters the vestibular region via a swelling level with, but continuous with, the ampulla for the posterior canal. The posterior semicircular canal (psc, Fig. 5E, G) is steeply inclined, almost vertical, and is the shortest of the three. Each canal terminates at a large, bulbous ampulla (aa, ha, pa, Fig. 5) roughly a third to a quarter of the length of each canal. All ampullae are confluent with the cranial cavity and fully enclosed in bone.

A sub-triangular utriculus (utr, Fig. 5E, F) lies immediately ventral to the anterior and horizontal ampullae. Both the posterior ampulla and utriculus join the deep, rounded saccular chamber (sac), which is only narrowly divided from its counterpart by the parachordal plates.

Both pairs of anterior and posterior semicircular canals meet at the crus commune (cc, Fig. 5G), which is narrowly separated from its antimere on the midline. The crus commune is inset into the dorsal surface of the endocast, ventral to the endocranial roof. A large lateral cranial canal (lcc, Fig. 5E. G) protrudes from the cranial cavity between the semicircular canals, although due to weak endochondral mineralisation, its posterior extent is poorly resolved. The lateral cranial canal is connected to the cranial cavity anteriorly and posteriorly through the loops of the semicircular canals and extends dorsally above the endocranial roof. It lacks a connection with either the posttemporal fossa or fossa bridgei.

### Cheek

The orbit is very large, extending almost a third of the length of the entire skull. The right circumorbital series is completely preserved, and consists of numerous bones: three supraorbitals, the dermosphenotic, two suborbitals (including an accessory suborbital, following the terminology of Arratia, 2013), five infraorbitals, and the antorbital. Only the second and third supraorbitals are present on the left side of the skull. Although the individual circumorbital bones are in almost continuous contact with each another, they do not form a complete ring, as supraorbital 1 (spo 1, Figs. 2B, D, 6B) and the antorbital (an, Figs. 2B, D, 6B) are interrupted by the nasal (na, Figs. 2B, D, 6B) and laterodermethmoid.

**Figure 6:**
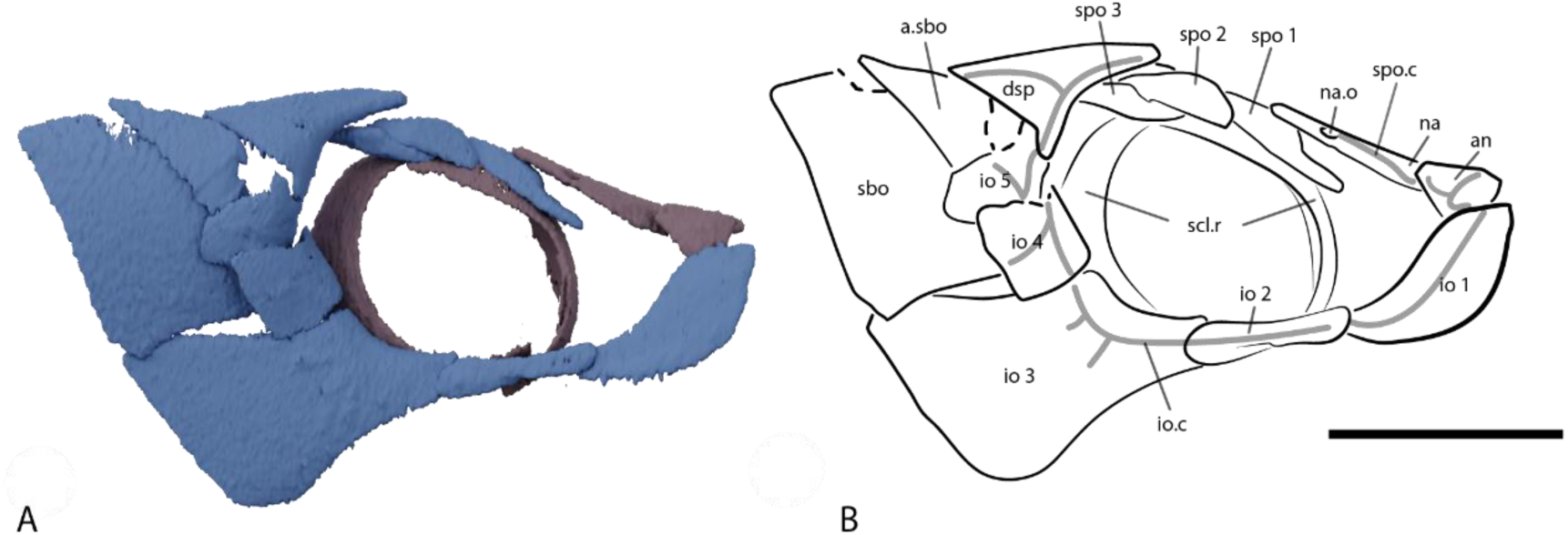
The circumorbital series of *Dorsetichthys bechei* NHMUK PV P 73995. (A) Render and (B) interpretive in right lateral view. Colour coding: blue, cheek; purple, skull roof. Scale bar = 10 mm. Abbreviations*: an, antorbital; dsp, dermosphenotic; io 1-5, infraorbital bones; io.c, infraorbital canal; na, nasal; na. o, nasal opening; sbo, suborbital; spo 1-3, supraorbital bones; spo.c, supraorbital canal*.

The three anamestic supraorbitals contact each other in sequence. Supraorbital 1 (spo 1, 2B, D, 6B) is elongate, with a narrow projection extending anteriorly beneath the nasal. Supraorbital 2 (spo 2, 2B, D, 6B) is broader, gently tapering posteriorly where it rests on top of supraorbital 3. Supraorbital 3 (spo 1, 2B, D, 6B) is irregularly shaped, with a small shelf-like overlap area to accommodate supraorbital 2.

The infraorbitals also form a continuous, uninterrupted series. Infraorbital 1 (io 1, Figs. 2B, 6B) is a large, smooth, curved plate which partially overlaps supramaxilla 1, the maxilla and the antorbital (an, Fig. 2B). Infraorbital 2 (io 2, Figs. 2B, 6B) is narrow and sub-rectangular, overlapping both neighbouring infraorbitals. Infraorbital 3 (io 3, Figs. 2B, 6B) is by far the largest of the circumorbital series, with a concave anterodorsal margin framing the posterior limit of the orbit. It broadens posteriorly into a roughly rectangular plate, partially overlapping the hyomandibula, and its anterior margin is embayed for articulation with infraorbital 2. Infraorbitals 4 and 5 (io 4, 5, Figs. 2B, 6B) are roughly square and almost equal in size, with narrow projections at their anterodorsal margins. Infraorbital 4 overlaps a significant portion of infraorbital 5. The infraorbital sensory canal (io.cl, Fig. 6B) is transmitted through each infraorbital, with at least three posterior ramifying branches in the third infraorbital.

Two suborbital (sbo, Figs. 2B, 6B) bones are present, partially overlapped by the infraorbitals. The smaller, accessory suborbital is triangular, with a straight dorsal margin roughly parallel with the edge of the skull roof. It overlaps the much larger, ventral suborbital, which forms a tall, sub-rectangular plate with an angled dorsal margin.

The dermosphenotic (dsp, Figs. 2B, D, 6B) is triangular, tapering to rounded points anteriorly, posteriorly, and ventrally. It has been slightly displaced to project above the supraorbitals and skull roof, but its short ventral ramus remains in contact with the fifth infraorbital. It houses the junction of the infraorbital canal between the infraorbital series and skull roof. An anterior branch of the canal is also transmitted above the orbit.

Finally, the sclerotic ring (scl.r, Figs. 2B, 6B) is formed of two ossicles, both of which are completely preserved on the right side of the skull. They contact dorsally but are offset by a gap ventrally, failing to form a continuous ring. Only displaced fragments of the left sclerotic ring are preserved.

### Shoulder girdle

The shoulder girdle is incomplete, with only the right extrascapular and posttemporal completely preserved within the matrix. The cleithrum is difficult to resolve in the CT data and not fully visible on the surface of the specimen.

The posterior margin of the skull roof is overlapped by a single large, triangular extrascapular (exc, Figs. 2B, D). It is thin and narrows considerably towards the midline, with the thicker lateral margin carrying the lateral line canal and at least five short laterally directed branches. There is a distinct medial commissure of the lateral line canal within the extrascapular. The extrascapular appears to extend beyond the lateral margin of the skull roof, creating an overhang above the operculum. A large, strongly curved posttemporal lies posterior to the extrascapular, although it has been displaced somewhat ventrally. Its anterior margin is concave to match the convex posterior margin of the extrascapular. The posttemporal has a thickened ventral lateral margin to accommodate the lateral line canal, and several small branches extend medially from the main trunk to open in pores on the surface.

### Upper jaw

The articulated right upper jaw includes the premaxilla, maxilla and two supramaxillae, while the left side preserves only a fragmentary premaxilla, displaced maxilla and the remnants of one supramaxilla (Figs. 2, 7, 8).

**Figure 7:**
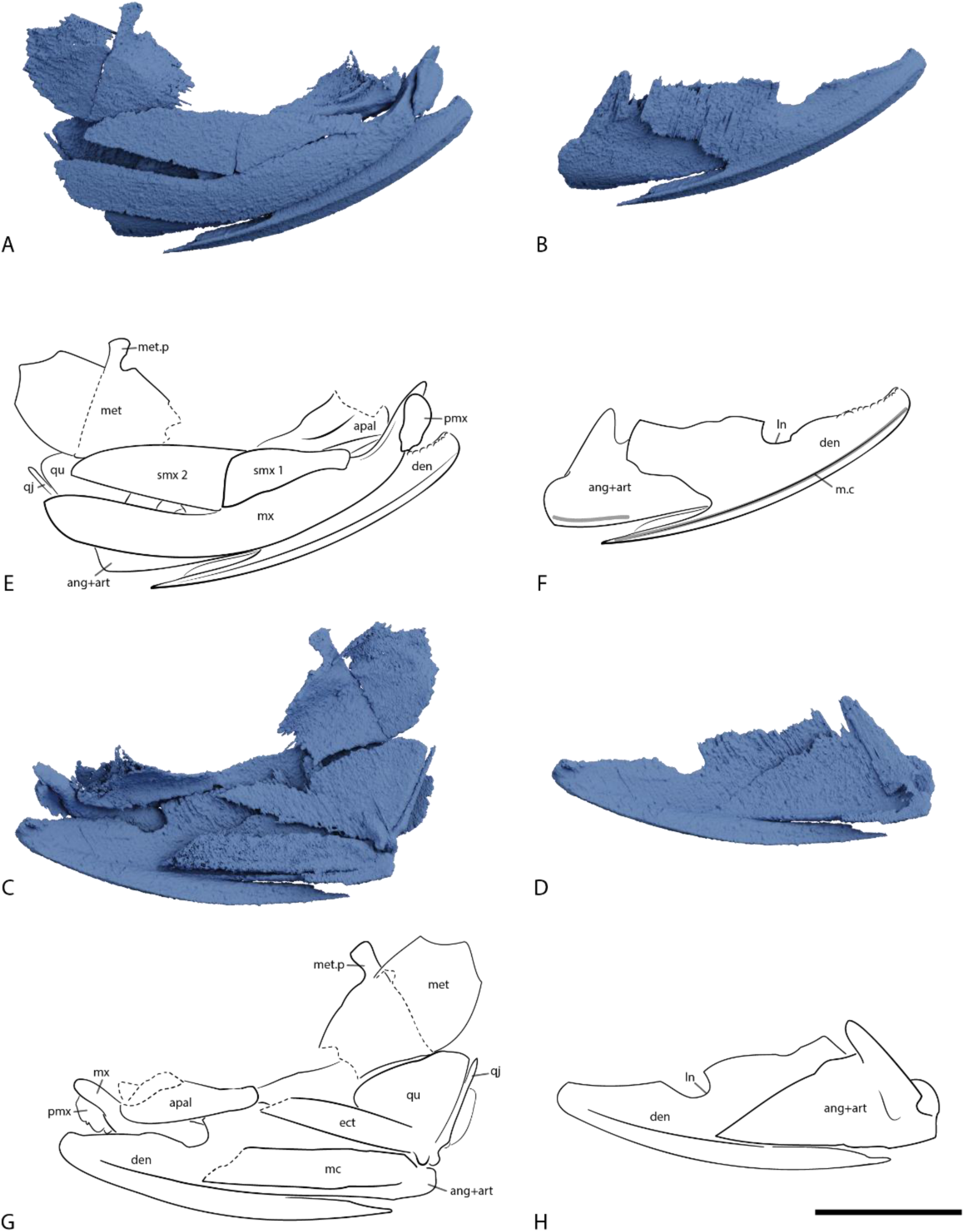
The right upper and lower jaws of *Dorsetichthys bechei* NHMUK PV P 73995. (A) Render and (E) interpretive drawing in right lateral view. (B) Render and (F) interpretive drawing in right lateral view, with upper jaw and palatoquadrate removed. (C) Render and (G) interpretive drawing in left lateral view. (D) Render and (H) interpretive drawing in left lateral view, with upper jaw and palatoquadrate removed. Scale bar = 10 mm. Abbreviations*: apal, autopalatine; ang+art, fused angular and articular; den, dentary; ect, ectopterygoid; ln, ‘leptolepid notch’; mc, Meckel’s cartilage m.c, mandibular canal; met, metapterygoid; met.p, metapterygoid process; mx, maxilla; pmx, premaxilla; smx, supramaxilla, qj, quadratojugal process; qu, quadrate*.

**Figure 8:**
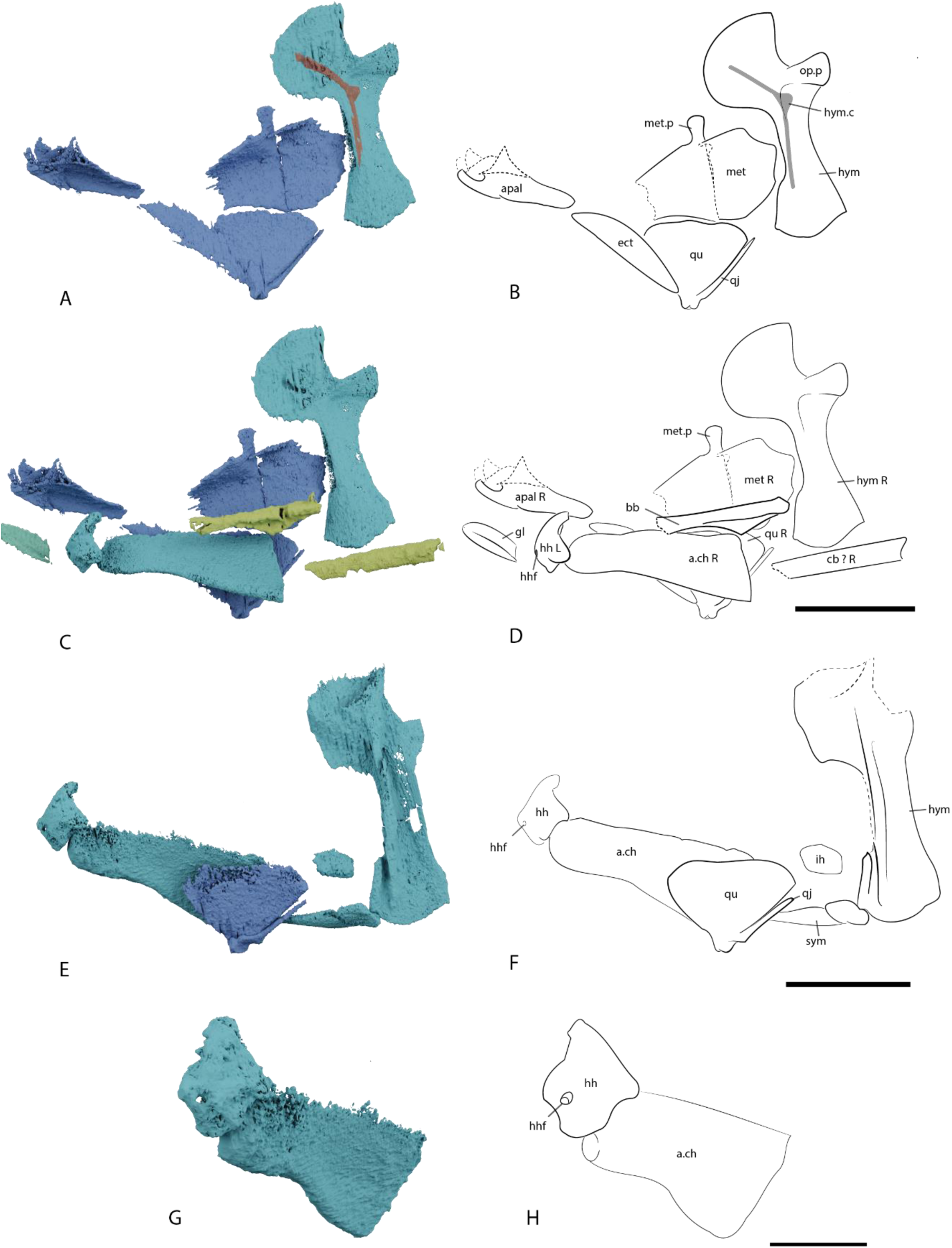
The palatoquadrate and hyoid arch of *Dorsetichthys bechei* NHMUK PV P 73995. (A) Render and (B) interpretive drawing of right palatoquadrate and hyoid arch in right lateral view. (C) Render and (D) interpretive drawing of the palatoquadrate, hyoid arch and gill skeleton in right lateral view, with. (E) Render and (F) interpretive drawing of left palatoquadrate and hyoid arch in right lateral view. Scale bar = 10 mm. (G) Render and (H) interpretive drawing of right hypohyal and anterior ceratohyal in anteroventral view Scale bar = 5 mm. Colour coding: blue, cheek and jaw; turquoise, hyoid arch; yellow, gill skeleton. Abbreviations: *apal, autopalatine; bb, basibranchial; cb, ceratobranchial; a.ch, anterior ceratohyal; ect, ectopterygoid; gl, gular plate; hh, hypohyal; hhf, efferent hyoidean artery; hym, hyomandibular, hym.c, hyomandibular canal; ih, interhyal; met, metapterygoid; met.p, metapterygoid process; op.p opercular process; qj, quadratojugal process; qu, quadrate; sym, symplectic*.

The premaxilla (pmx, Figs. 4B, D, 7A, 8G) is small and sub-triangular, bearing a single row of approximately six small teeth. Each tooth is small, blunt and peg-like, and directed medially. The left premaxilla bears a short premaxillary process (pmx.p, Fig. 4B) which articulates with a small socket on the anterior face of the lateral dermethmoid (amx, Fig. 4D), indicating that the premaxillae were capable of rotational movement. Its inner surface bears a ridge, above which rests the peg-like anterior process of the maxilla.

The maxilla (mx, Fig. 7E, G) is gently curved, elongate and substantially overlaps the mandible over its entire length. A single row of minute teeth appears to be present along the ventral margin of the bone. These are at the limit of scan resolution and their number and geometry cannot be accurately discerned. The outer surface bears ridged ornament that is subparallel to the long axis of the bone, while the inner surface is relatively smooth, although the dorsal half of the bone is thicker than the ventral half. Anteriorly, the maxilla narrows and projects towards the midline and somewhat dorsally behind the premaxilla. The anterior process of the maxilla forms a narrow peg that articulates with the ethmoid, terminating in a foramen in the lateral dermethmoid (fet, Fig. 4K).

Two thin supramaxillae are preserved on the right side of the specimen. The posterior supramaxilla (smx 2, Fig. 7E, G) is twice as large as the anterior supramaxilla (smx 1, Fig. 7E, G). Supramaxilla 1 is sub-rectangular and tapers anteriorly, partially overlapping the maxilla, while supramaxilla 2 is more elongate, with a straight ventral margin and a curved dorsal margin, tapering to a point posteriorly. A narrow dorsal projection extends beneath supramaxilla 1. A gap between supramaxilla 2 and the maxilla suggests with that the supramaxilla is displaced or its ventral margin is too thin to be resolved in the scan.

### Palate

The palate includes the quadrate, metapterygoid, ectopterygoid, entopterygoid and autopalatine. The quadrate (qu, Figs. 7E, G, 8B, D, F) is broadly triangular with a thickened, well ossified posteroventral margin, developed as two cotyles to articulate with the lower jaw. A long, splint-like process, sometimes referred to as the quadratojugal process (qj, Figs. 7E, G, 8B, D, F), projects posterodorsally from the body of the quadrate. The process lies parallel to the posterior margin of the bone, and tapers to a point.

The large metapterygoid (met, Figs. 7E, G, 8B, D) lies dorsal to the quadrate, with an anterior face that curves medially and bears a well-developed metapterygoid process (met.p, Figs. 7E, G, 8B, D). It is incompletely ossified anteriorly. The ectopterygoid (ect, Fig. 9B) is elongate and rectangular. It does not appear to be tooth-bearing, is extremely reduced, and projects anterodorsally from the jaw joint. The entopterygoid, like the anterior portion of the metapterygoid, is too weakly mineralised to be accurately segmented. However, examination of the tomograms suggests it fills the gap between the metapterygoid and the autopalatine.

**Figure 9:**
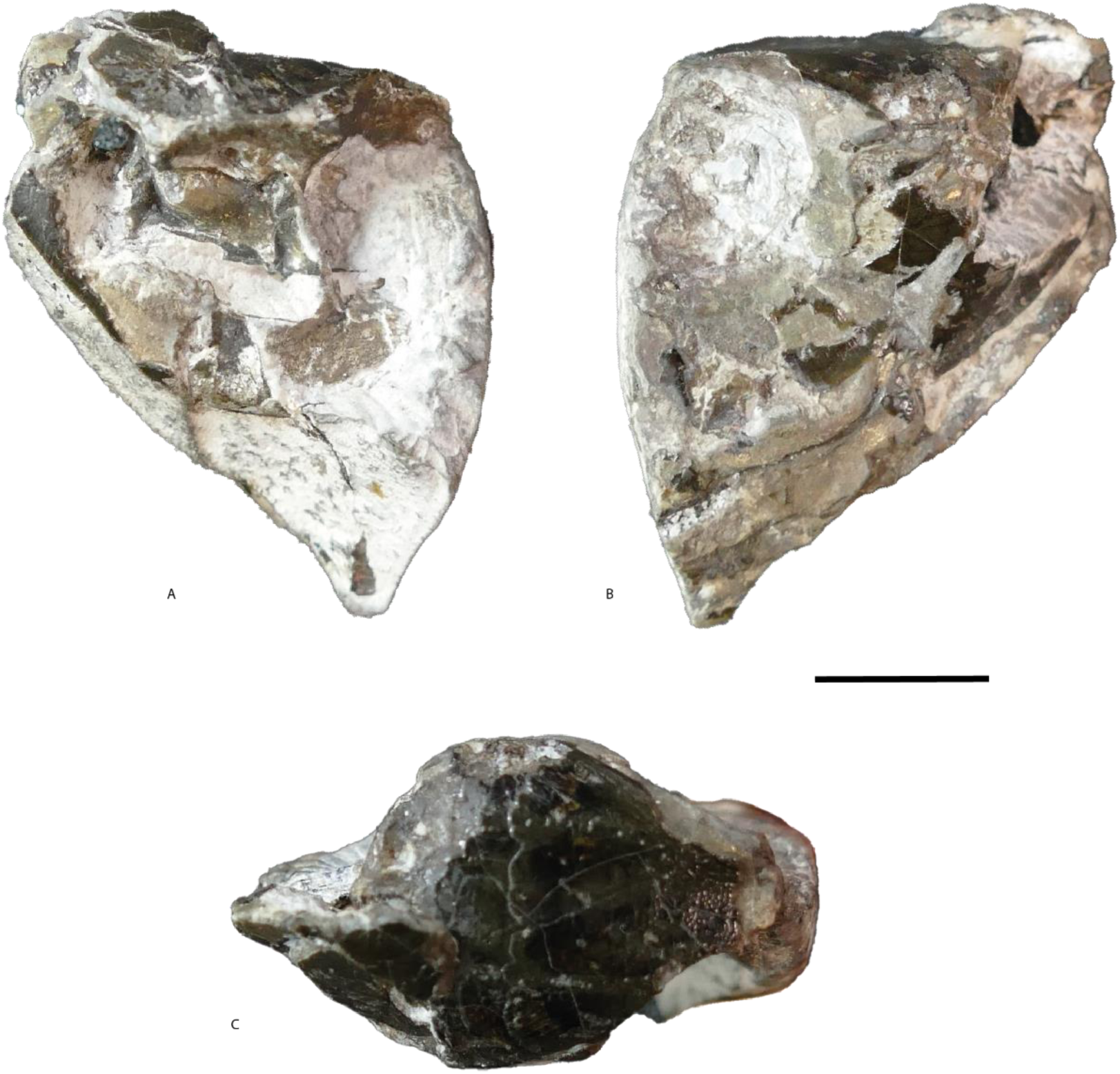
*Dorsetichthys bechei* NHMUK PV P 1052 in (A) right lateral, (B) left lateral and (C) dorsal view. Scale = 10 mm.

The autopalatine (apal, Figs. 7E, G, 8B, D) is robustly ossified and tightly bound fused to the lateral ethmoid; together, these floor the anterior corner of the orbit. Its dorsal surface is smooth and gently concave, with a thickened lateral margin.

### Lower jaw

The lower jaw (Fig. 7) consists of the dentary, Meckel’s cartilage, and a fused angular and articular. Both the left and right dentaries (den, Fig. 7) are well preserved, although slightly out of alignment. The dentary is curved and elongate, with an anterior portion that curves strongly towards the midline. A deeply embayed posterior margin of the dentary accommodates the angular. Pyrite lines the dorsal margin of the dentary, making the presence or absence of teeth difficult to determine. There may be some teeth close to the anterodorsal margin (Fig. 7E, F). Approximately halfway along the dorsal margin of the dentary, which slopes anteroventrally, is a wide, sub-rounded indentation: the so-called “leptolepid notch” (ln, Fig. 7F, H). Both the outer and inner surface of the dentary are smooth. Ventrally, the dentary thickens into a lateral ridge, with a distinct ventromedially directed flange. The mandibular canal (m.c, Fig. 7F) is housed within the bone at the junction of this ridge and flange. It extends from the anterodorsal tip of the dentary to its posterior margin, after which it passes into the angular. At least half a dozen small, equally spaced pores, infilled with pyrite, connect the mandibular canal to the external surface of the dentary.

The large, triangular angular is significantly overlapped by the dentary, with its anterior apex reaching to the level of the “leptolepid notch”. A thickened ridge along the ventral margin of the angular houses the mandibular canal. Its dorsal margin is approximately level with that of the dentary, and together these form a modest coronoid process. There is no independent surangular. At its posteroventral corner, the angular is fused to the articular (ang+art, Fig. 7); the posterior margin of this compound ossification bears two articulation facets to receive the condyles of the quadrate. Meckel’s cartilage (mc, Fig. 7G) is ossified anterior of the articular to the level of the “leptolepid notch”. It is rectangular and lies parallel to the dentary, with slightly thickened dorsal and ventral margins. Although its anterior extent is difficult to determine due to poor ossification, there is no indication of an independent anterior mentomeckelian ossification. A thin, rectangular element that is challenging to segment lies closely applied and parallel to the upper portion of Meckel’s cartilage on both jaws. Comparison to the better-resolved lower jaws of NHMUK PV P 1052 indicates that this ossification is a prearticular. It does not appear to be tooth-bearing, but this may be a limit of the scan resolution. Coronoids are absent. The posteroventral corner of the lower jaw barely extends posterior to the articular facets, and there is no evidence of a well-developed postarticular process or retroarticular.

#### Hyoid Arch

The hyoid arch comprises the hyomandibula, ceratohyal, hypohyal, interhyal and symplectic. Both the right (hym, Fig. 8B, D) and left hyomandibula (hym, Fig. 8F) are preserved, but the right is significantly better preserved and *in situ*, in articulation with the hyomandibular facet on the braincase. The hyomandibula is broad, with an upright ventral limb and large anterior head that lies at an acute angle to the rest of the ossification. Both the ventral and anterodorsal margins are convex, and a stout opercular process is present on the posterior margin. A pronounced ridge, which flares dorsally, runs down the lateral face of the hyomandibula. A corresponding, although much fainter, ridge is present on the medial face. The hyomandibular nerve (hym.c, Fig 8B) pierces the medial face of the hyomandibula anterodorsal to the conjunction of the vertical ridge and opercular process, exiting the lateral face posteroventrally and continuing in a groove on the posterior margin of the ridge.

The symplectic (sym, Fig. 8F) is elongate, extending medial to the quadrate. It articulates with the hyomandibula posteriorly and is not involved in the lower jaw joint. Only the anterior ceratohyal (a.ch, Fig. 8D, F, H) is preserved; it is likely that the posterior ceratohyal was not ossified and was instead cartilaginous. Both the left and right anterior ceratohyal are well preserved, although the right ceratohyal is significantly displaced and overturned. It has a thickened, rounded ventral margin, but is poorly ossified along its dorsal margin. Unusually, the ceratohyal lacks a groove for the afferent hyoidean artery. The hypohyal (hh, Fig. 8D, F, H) also forms a single ossification on each side, as opposed to dorsal and ventral ossifications. It is very weakly ossified and is pierced by the efferent hyoidean artery via a foramen on the ventral margin (hhf, Fig. 8D, F, H). The right interhyal (ih, Fig. 8F) is also preserved, though it is displaced dorsally.

### Gill skeleton

The gill skeleton is very poorly mineralised in this specimen; several elements appear faintly in tomograms, but only a basibranchial and fragment of a ceratobranchial can be segmented. The basibranchial (bb, Fig. 8D) lies directly ventral and parallel to the parasphenoid. It is elongate, with a shallow, triangular ridge along its ventral margin and a flared anterior margin. The ceratobranchial (cb, Fig. 8D) is present ventrolateral to the braincase. It is narrow, with a U-shaped trough along its ventral margin. Other bones posterior to the braincase might also represent fragments of gill skeleton, but they are too faintly ossified to be accurately segmented.

### Operculogular series

Elements of both the left and right opercular series are present. Only the opercle (op, Fig. 2B) can be resolved in tomograms, although other elements are visible externally. The left preopercular is expanded ventrally and narrows dorsally. Its anterior margins are overlapped by the suborbitals. The path of the preopercular canal and any associated rami is not externally visible. The opercle forms a large, ovoid plate, the anterior margin of which appears to abut the preopercle. The posterior margin of the opercle is well-rounded. No other elements of the opercular skeleton can be described. A thin, ovoid median gular plate (gl, Fig. 8D) appears to be present, lying between the left and right dentaries.

## Description: NHMUK PV P 1052

### General morphology

NHMUK PV P 1052 (Fig. 9) includes an almost complete braincase lacking only the ethmoid region, as well as partially preserved elements of the skull roof, gill skeleton, cheek, circumorbital series, palate, and lower jaw, all partly encased in matrix. The anterior portion of the specimen, beyond the orbit, is missing. Many dermal plates have been also displaced or lost.

The anatomy of NHMUK PV P 1052 is largely similar to that of NHMUK PV P 73995, including the braincase and skull roof (Figs. 10, 11). As such, we do not redescribe these elements here, and instead focus on differences in anatomy or features that cannot be described in NHMUK PV P 73995. Furthermore, although the braincase of NHMUK PV P 1052 is more complete, its internal endocranial walls are poorly mineralised, and the endocast is difficult to segment accurately (Fig. 12) and it is therefore not described in detail.

**Figure 10:**
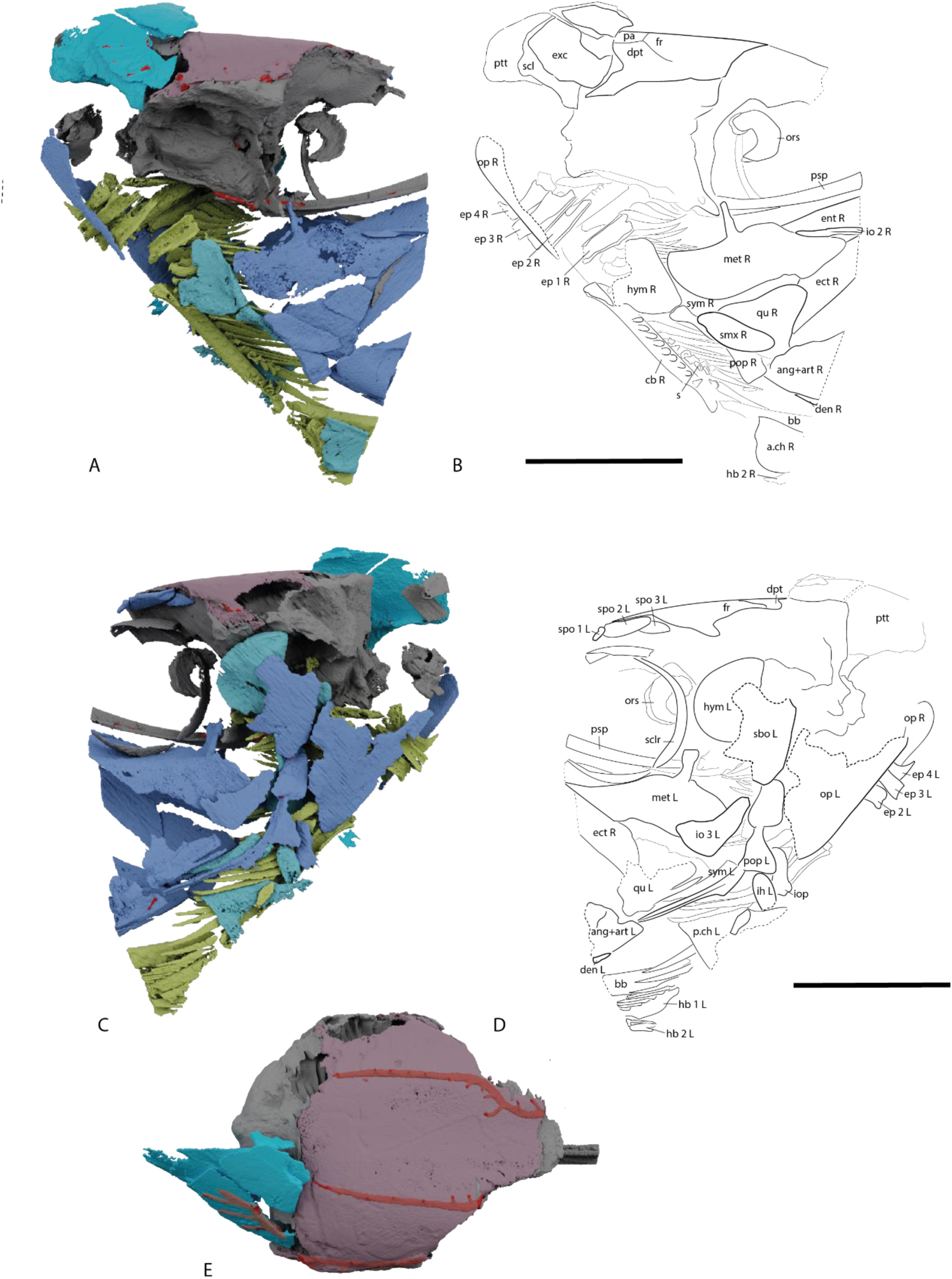
*Dorsetichthys bechei* NHMUK PV P 1052. (A) render and (B) interpretive drawing in right lateral view, (C) render and (D) interpretive drawing in left lateral view, render (E) in dorsal view. Colour coding: blue, cheek and jaw; purple, skull roof, dermal shoulder girdle and sclerotic ossicle; grey, braincase; turquoise, hyomandibula; light green, operculogular system; yellow, gill skeleton. Sensory lines in grey Scale = 10 mm. Abbreviations: *ang+art, fused angular and articular; bb, basibranchial; a.ch, anterior ceratohyal; p.ch, posterior ceratohyal; den, dentary; dpt, dermopterotic; ect, ectopterygoid; ent, entopterygoid; exc, extrascapular; fr, frontal; hb, hypobranchial, hym, hyomandibular; ih, interhyal; io, infraorbital; iop, interoperculum; met, metapterygoid; op, opercular; ors, orbitosphenoid; pa, parietal; pop, preopercular; psp, parasphenoid; ptt, posttemporal; qu, quadrate; s, gill raker posterior socket; scl, sclerotic ring; smx, supramaxilla; spo, supraorbital; sym, symplectic*.

**Figure 11:**
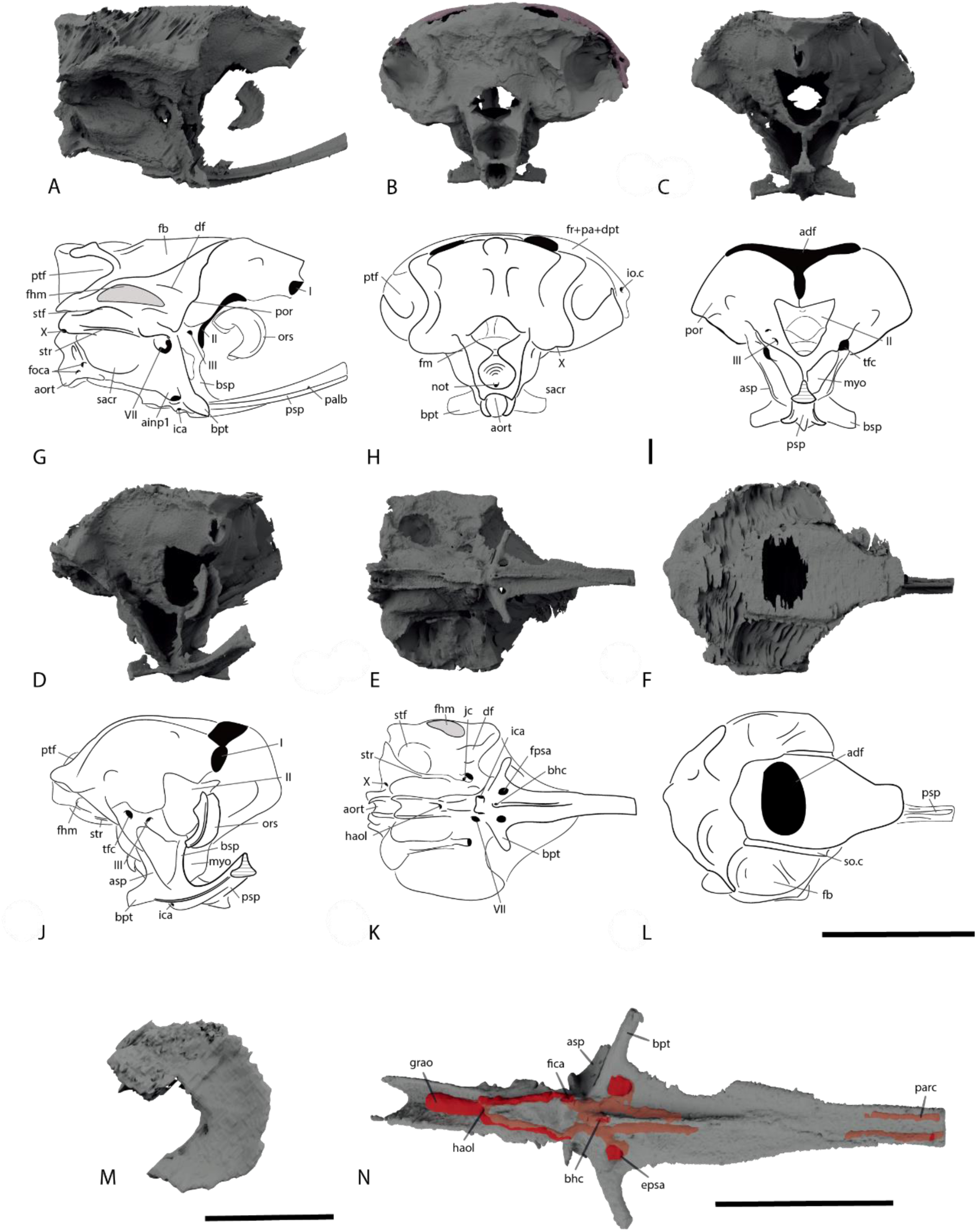
The braincase of *Dorsetichthys bechei* NHMUK PV P 1052. Renders in (A) right lateral, (B) posterior, (C) anterior, (D) anterolateral, (E) ventral and (F) dorsal view. Interpretive drawings in (G) right lateral, (H) posterior, (I) anterior, (J) anterolateral, (K) ventral and (L) dorsal view. Scale bar = 10 mm. (M) Render of orbitosphenoid in right lateral view. Scale bar = 2.5 mm (N) Render of parasphenoid in ventral view. Scale bar = 5 mm Orbitosphenoid not rendered in panels (B-L). Abbreviations*: adf, anterior dorsal fontanelle; aort, aortic notch; bhc, buccohypophysial canal; bpt, basipterygoid process; bsp, basisphenoid; df, dilatator fossa; epsa, efferent pseudobranchial artery; fb, fossa bridgei; fhm, hyomandibular facet; fm, foramen magnum; fotc, otoccipital fissure; fr+pa+dpt, fused frontal, parietal and dermopterotic (skull roof); haol, housing of aortic ligament; ica, internal carotid artery; myo, posterior myodome; not, notochord; pf, pituitary fossa; por, post-orbital process; psp, parasphenoid; ptf, post-temporal fossa; sacr, saccular recess; socc, supraoccipital; spic, spiracular canal; str, lateral strut; tfr, trigemino-facial recess; II, optic fenestra; VI, foramen of abducens nerve; VII, foramen of facial nerve X, foramen of vagus nerve*.

**Figure 12:**
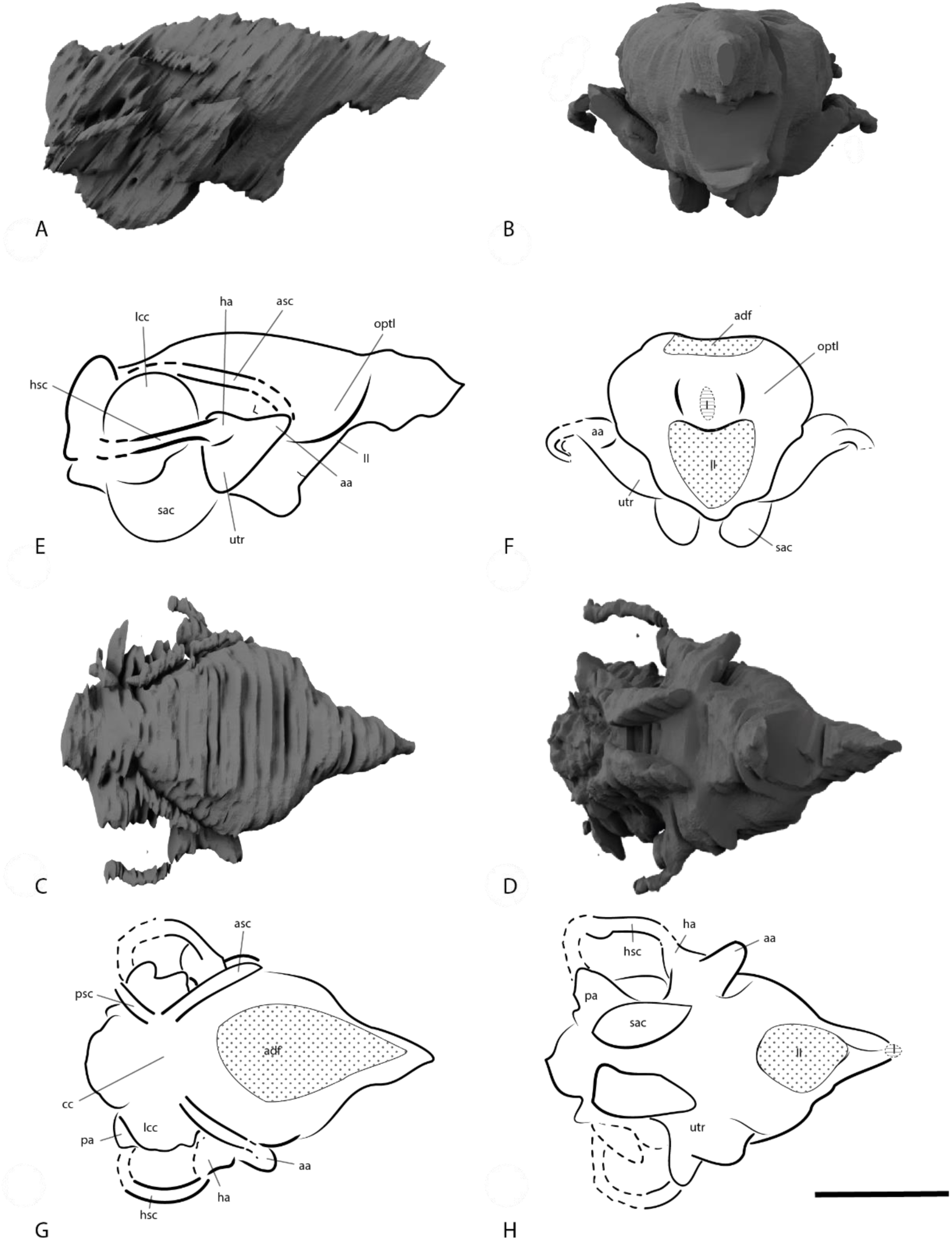
The endocast of *Dorsetichthys bechei* NHMUK PV P 1052. Renders in (A) right lateral, (B) anterior, (C) dorsal and (D) ventral view. Interpretive drawings in (E) right lateral, (F) anterior, (G) dorsal and (H) ventral view. Scale bar = 5 mm. Abbreviations*: aa, anterior ampulla; asc, anterior semicircular canal; aur, cerebellar auricles cc, crus commune; ha, horizontal ampulla; hsc, horizontal semicircular canal; ica, internal carotid artery; lcc, lateral cranial canal; not, notochordal canal; optl, optic lobe; psc, posterior semicircular canal; sac, saccular; utr, utricular; II, optic nerve; VI, abducens nerve; X, vagus nerve*.

### Braincase

The braincase of this specimen has already been described by Rayner (1948) and Patterson (1975) on the basis of external examination, and we have little to add to their comprehensive descriptions. Most of the braincase is ossified as a single unit comprising the occipital, otic, orbitotemporal and basioccipital regions, with the exception of the separate orbitosphenoid. The ethmoid region is not preserved as the specimen is incomplete anteriorly. Grooves on the posterior face of the occiput have been interpreted as delimiting the epioccipitals, exoccipitals and supraoccipital (Patterson 1975: fig. 55), but these cannot be traced as separate ossifications extending into the neurocranium in tomograms.

In most aspects, the braincase is identical to that of NHMUK PV P 73995. The separate orbitosphenoid ossification is well mineralised, and can be described in detail (Fig. 11M). It resembles a backwards ‘C’, and occupies less than a quarter of the orbit. Although the orbitosphenoid forms a single median ossification, it appears to comprise two plates that join posteriorly and diverge anterolaterally. A deep notch in the posterior margin opposes the opening of the optic nerves into the orbit.

### Parasphenoid

The parasphenoid (Fig. 11N) is very well preserved, allowing the route of the basicranial circulation to be reconstructed. The parasphenoid is applied directly to the underside of the basiocciput, with the convex form of the posterior stalk of the parasphenoid fitting into a corresponding concave surface on the base of the braincase. The dorsal aorta extended along the deep aortic groove (grao, Fig. 11N) which terminates at the median, cup-like housing of the insertion of the aortic ligament (haol, Fig. 11N). The dorsal aorta bifurcates around the opening, continuing in shallow grooves on the lateral face of the parasphenoid before entering laterally directed foramina for the internal carotid arteries (ic, Fig. 11N). The foramina for the internal carotids are housed in a ventral pedestal. The internal carotids emerge on the dorsal surface of the parasphenoid and briefly run along shallow grooves before bifurcating once again. One branch flows laterally and enters the wide, paired openings of the efferent pseudobranchial arteries (epsa, Fig. 11N), while the other continues anteriorly along the median crest of the parasphenoid, following the parabasal canal (parc, Fig. 11N). The median, ventral opening of the buccohypophysial canal (bhc, Fig. 11N) is also present and positioned directly between the openings of the left and right foramina of the efferent pseudobranchial arteries. It has a rounded, well ossified rim and forms part of the ventral keel of the anterior corpus of the parasphenoid. As with NHMUK PV P 73995, teeth are not visible, but this may be an artefact of scan resolution.

### Cheek

The cheek is incomplete. Most dermal plates are disarticulated or heavily prepared and thus indistinguishable in tomograms, with only the preoperculum, one suborbital, two infraorbitals and two or three supraorbitals present. Other, smaller fragments of dermal bone cannot be identified.

One infraorbital (io 2, Fig. 10B, D) is present on the right side. It is very well preserved and morphologically identical to the second infraorbital of NHMUK PV P 73995. However, it has been displaced ventrally to lie on the palate. A fragment of infraorbital 3 can also be identified on the left side of the specimen (io 3, Fig. 10B). A large, plate-like suborbital (sbo, Fig. 10D) is present on the left side of the specimen and can be identified by its position relative to the orbit and the absence of a canal. At least two complete supraorbital bones (spo 1-3, Fig. 10D), identical to those of NHMUK PV P 73995, are preserved *in situ* on the left side. A small fragment of bone anterior to the two complete supraorbitals likely represents a part of the first supraorbital, giving NHMUK PV P 1052 a total of three supraorbitals matching NHMUK PV P 73995.

### Operculogular series

A partial preoperculum (pop L, Fig. 10D) can be identified on the left side of the specimen. Its posterior margin is difficult to discern, and the preopercular sensory canal, which extends along the anterodorsal margin, occupies the most complete portion. On the right side, only the rounded anteroposterior margin of the preoperculum is preserved within the matrix (pop R, Fig. 10B). The preopercular canal is transmitted through this into the back of the lower jaw, which it abuts. Similarly, only a portion of the large, left operculum (op, Fig. 10B, D) can be identified. Several short, posteriorly-projecting filaments line the medial face of the operculum. A very small portion of the interoperculum (iop, Fig. 10D) is preserved posterior to the preoperculum, but it is truncated by the specimen break, and little can be said of its morphology.

### Upper jaw

The dermal portions of the upper jaw are largely missing, with only one supramaxilla preserved on the right side, lateral to the quadrate (smx R, Fig 10B). It is lenticular in shape and exposed at the surface, where the specimen is incomplete posteriorly.

### Palate

The palate is very well ossified and preserved in articulation, comprising the metapterygoid, entopterygoid, ectopterygoid and quadrate. Its morphology largely matches that of NHMUK PV P 73995. The metapterygoid (met, Figs. 10B, D, 13B, D, E) is a large, sheet-like ossification with a rectangular lateral face and a triangular medial face that extends towards the parasphenoid. This medial face tapers anteriorly, substantially overlapping the posterior portion of the entopterygoid and, to a lesser extent, the ectopterygoid. A well-developed metapterygoid process (met.p, Figs. 10B, D, 13B, D, E) extends dorsally from the junction between these faces to articulate with the anterior margin of the basipterygoid process. The preserved portion of the entopterygoid (ent, Figs. 10B, D, 13B, D) is large and ovoid, and contacts its antimere on the midline anteriorly, excluding most of the parasphenoid from the oral cavity except for the posterior portion of the narrow ventral ridge. As in NHMUK PV P 73995, the quadrate (qu, Figs. 10B, D, 13B, D, E) is sub-triangular, with an elongate splint-like posterodorsal process. Anteroventrally, the quadrate is expanded into condyles that articulate with the lower jaw. The palate appears to be edentulous, with smooth medial surfaces.

**Figure 13:**
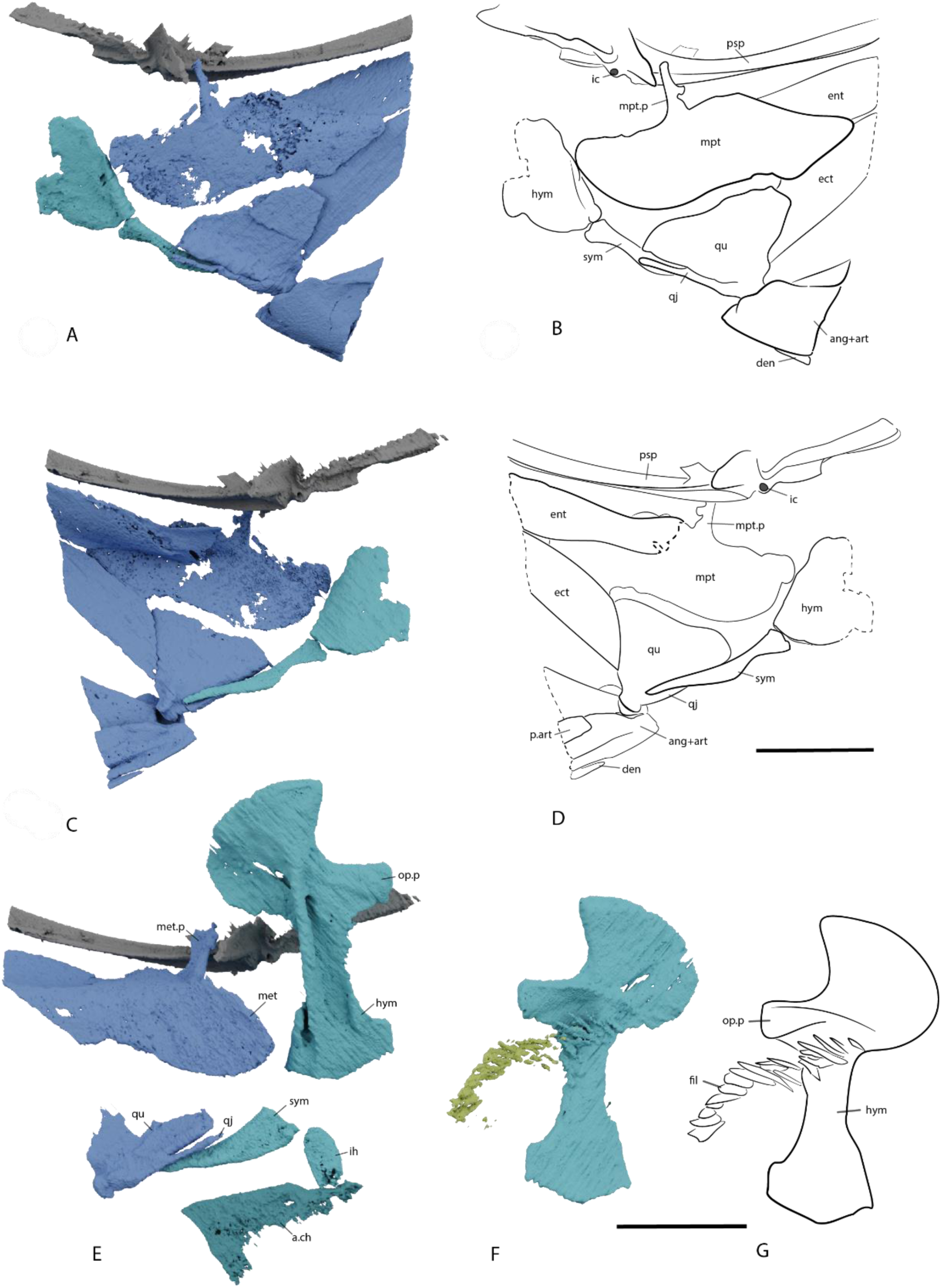
The palatoquadrate and hyoid arch of *Dorsetichthys bechei* NHMUK PV P 1052. (A) Render and (B) interpretive drawing of right palatoquadrate and hyoid arch in right lateral view. (C) Render and (D) interpretive drawing of right palatoquadrate and hyoid arch in left lateral view. (E) Render of left palatoquadrate and hyoid arch in left lateral view. (F) Render and (G) interpretive drawing of left hyomandibular and associated gill filaments. Colour coding: blue, cheek and jaw; turquoise, hyomandibula. Scale bar = 5 mm. Abbreviations*: ang+art, fused angular and articular; a.ch, anterior ceratohyal; den, dentary; ect, ectopterygoid; ent, entopterygoid; fil, fillaments; hym, hyomandibular; ic, internal carotid; ih, interhyal; met, metapterygoid; met.p, metapterygoid process; op.p, opercular process; p.art, prearticular; psp, parasphenoid; qj, quadratojugal process; qu, quadrate; sym, symplectic*.

### Lower jaw

A short portion of the angular and articular are preserved in articulation. The angular is narrow and carries the mandibular sensory canal along its ventral margin. The articular and angular are fused (ang+art, Figs. 10B, D, 13B, D), and the posterodorsal surface of the articular forms a deep articulation facet for the quadrate. Ossification of the Meckel’s cartilage continues anteriorly until the specimen break. A rectangular plate, which we identify as the prearticular (p.art, Fig. 13D), lies along the dorsal portion of the medial face of the mandible. Only a tiny fragment of the posteroventral margin of the dentary (den, Figs. 10B, D, 13B, D) is preserved.

### Hyoid arch

The hyoid arch includes a hyomandibula, ceratohyal and symplectic. The left interhyal is also preserved. The left hyomandibula (hym, Figs. 10D, 13E, F) is fully intact, while only a fragment of the right hyomandibula (hym, Fig. 13B, D) is present. Its gross morphology appears to match that of NHMUK PV P 73995. In addition, a series of sinuous, posteriorly directed filaments (gf, Fig. 13G) are present on the medial surface of the left hyomandibula, ventral to the opercular processes. They form a continuous series with the filaments borne on the posterodorsal margin of epibranchial 1, and may represent evidence of a pseudobranch, functioning alongside the spiracular canal (Laurent & Dunel-Erb, 1984).

Only a small portion of the right anterior ceratohyal (a.ch, Fig. 14D) is preserved due to a break in the specimen, positioned ventral to the braincase along with the ventral gill skeleton. It is narrow and roughly rectangular in axial section, with a thickened ventral margin. The left posterior ceratohyal (p.ch, Fig. 13E) is near complete, though exposed on the surface ventrally. It bears a broad notch on its dorsal surface to accommodate the interhyal (ih, Fig. 13G), and is not attached to any associated branchiostegals. The symplectic (sym, Fig. 13B, D, E) is identical to that of NHMUK PV P 73995. The right symplectic (sym, Fig. 13B, D, E) is preserved in articulation with the hyomandibula posteriorly. Both the left and right symplectic terminate anteriorly against the inner surface of the quadrate, lying parallel to the quadratojugal process (qj, Fig. 13B, D, E)

**Figure 14:**
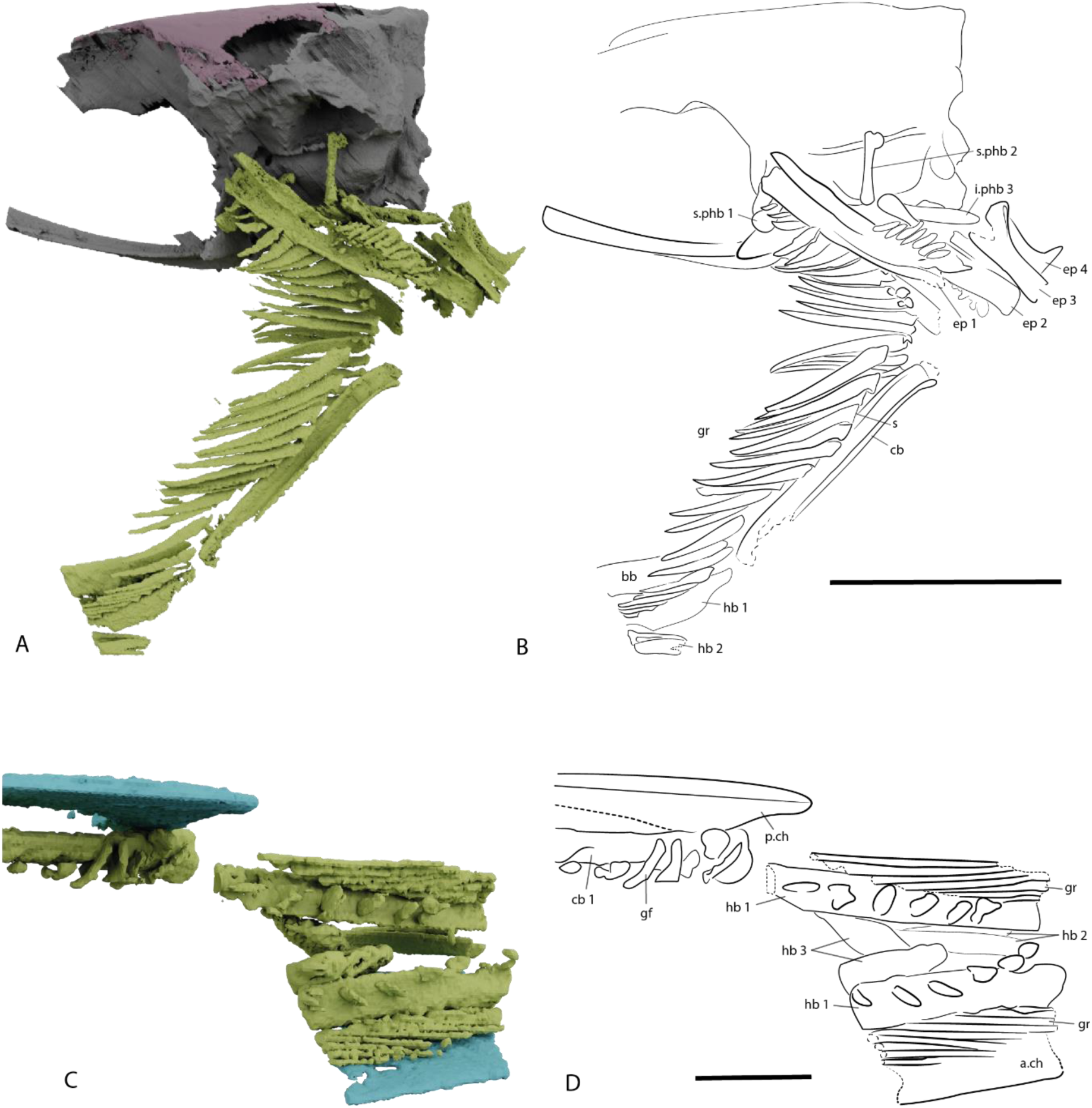
The gill skeleton of *Dorsetichthys bechei* NHMUK PV P 1052. (A) Render and (B) interpretive drawing of complete gill skeleton in left lateral view. Scale bar = 10 mm. (A) Render and (B) interpretive drawing of ventral gill skeleton and hyoid arch in dorsal view. Scale bar = 2.5 mm. Abbreviations: *bb, basibranchial; cb, ceratobranchial; a.ch, anterior ceratohyal; p.ch, posterior ceratohyal; ep 1-4, epibranchials; gf, gill filaments; gr, gill rakers; hb 1-3, hypobranchials; i.phb 1-3, infraphrayngobranchials, s, gill raker posterior socket; s.phb 1-2, suprapharyngobranchials*.

### Gill skeleton

Unlike in NHMUK PV P 73995, the gill skeleton of NHMUK PV P 1052 is well mineralised and articulated (Fig. 14), and thus can be described in detail. The ventral gill series is incomplete on both sides, but includes ceratobranchials, hypobranchials and a basibranchial ossification. The dorsal gill series includes four epibranchials, two pharyngobranchials in the first two arches, and an infrapharyngobranchial in the third arch. Due to the way the specimen is preserved, dorsal components on the right side of the specimen are incomplete anteriorly. An array of rakers and filaments are also present on both sides. Though they are slightly displaced, rakers appear to form an uninterrupted orderly sequence across the dorsal and ventral component of the gill arch.

Only a single basibranchial (bb, Fig. 14B) is present, although the specimen is incomplete anteriorly and posteriorly, and multiple basibranchials are present in NHMUK PV P 73995. It is broader anteriorly and narrows somewhat posteriorly, although the anterior margin is incomplete. The dorsal surface is smooth and flat, and the ventral surface is developed into a broad median ridge. Paired articulation facets are preserved on either side of the ventral ridge, which would have articulated with the hypobranchials in life.

Only a single ceratobranchial is preserved, and its length and position immediately medial to the hyomandibula suggests it belongs to the first arch (cb, Fig. 14). It is complete on the left side of the specimen, but only a small fragment is present on the right. The ceratobranchial is elongate, with a slight medial deflection at the distal end, and a groove running along its entire ventrolateral surface. Its dorsal margin bears a row of at least eleven elliptical nodes, angled obliquely to the long axis of the cerotobranchial. 13 anteriorly-directed gill rakers (gr, Fig. 14B) are preserved in articulation with the ceratobranchial, and these rakers articulate with the lateral margin of the ceratobranchial nodes. Each raker is elongate and triangular, tapering anteriorly, with a concave dorsal margin and convex ventral margin. The posterior margin is thickened into a shallow, cup-shaped socket (s, Fig. 4B) for articulation with a corresponding node on the host ceratobranchial. In addition to nodes and rakers, the ventral quarter of the ceratobranchial bears a row of fine, undulous gill filaments, positioned medial to the nodes. Although the right ceratobranchial is mostly missing, some of its associated rakers are still present. Rakers of the ventral gill arch are largest nearer the posterior end of the ceratobranchial, and each overlaps the next in sequence slightly at its proximal end.

Three pairs of hypobranchials (hb, Fig. 14) articulate with the ventral face of the basibranchial, although the arch to which they belong cannot be determined with confidence. The more anterior pair are elongate, straight, and C-shaped in cross section, with a groove along the ventral margin, although are truncated anteriorly and posteriorly where the specimen is broken. The left aligns with the complete ceratobranchial, and therefore may represent hypobranchial 1 (hb1; Fig. 14). A series of elongate rakers, continuous with those of the ceratobranchial, articulate with the lateral surface of the hypobranchial, and short, stubby filaments are borne along its entire dorsal surface. A second pair of ossifications, preserved somewhat ventral to the others, resemble shorter, slightly flattened versions of the first pair, and may belong to the second gill arch (hb2; Fig. 14). The most posterior pair are fully enclosed within matrix and thus complete. They are short and gently curved, with no distinct groove, and are tentatively identified as hypobranchial 3 (hb3; Fig. 14). Rakers and filaments appears to be absent from the putative second and third hypobranchials.

Four epibranchials (ep 1-4, Fig. 14B) are preserved on each side. Epibranchial 1 is the largest of the four, and shows slight medial curve toward its distal end. Its dorsolateral face is grooved, and the anterodorsal corner is developed as a modest uncinate process. At least 14 gill rakers articulate with its medioventral margin via small nodes. The rakers are morphologically similar to those found on the ceratobranchial, although somewhat shorter, tapering anteriorly. At least twelve short, stout, straight elements in contact with the posterodorsal and posterior margin of the epibranchial likely represent gill filaments. Infrapharyngobranchial 1 and suprapharyngobranchial 1 are short, stout elements that near the anterior margin of the epibranchial; the infrapharyngobranchial is curved and the suprapharyngobranchial straight. Epibranchial 2 is only slightly shorter than the first, but is otherwise morphologically similar, also supporting two pharyngobranchials. Infrapharyngobranchial 2 (i.phb2, Fig. 14B) is around a third of the length of the epibranchial, straight, and without a groove, but bifurcates anteriorly to form a short dorsally directed process. Suprapharyngobranchial 2 (s.phb2, Fig 14 is narrow and rod-like. Approximately five short rakers project anterolaterally from the lateral face of epibranchial 2, and around eight very short filaments sit on its dorsomedial face, with a smaller number arranged in a ventromedial series. Epibranchials 3 and 4 are much shorter, straight, and do not seem to have associated rakers or filaments. The third epibranchial is U-shaped in cross section, and bears an uncinate process; the process on epibranchial 3 is proportionately the largest of all. Only one pharyngobranchial element is associated with this arch. Infrapharyngobranchial 3 is elongate, approaching the length of the second epibranchial, and laterally compressed. It also bifurcates anteriorly, and the anterior process is longer than the dorsal. Epibranchial 4 lacks a groove, has a very small uncinate process, and bears a short dorsal projection at its posterior end. A series of minute, scattered elements are present around the posterior portions of epibranchials 3 and 4, but it is impossible to tell whether these are rakers or filaments. The arrangement of the epibranchials and pharyngobranchial in life position is estimated in Figure 15.

**Figure 15:**
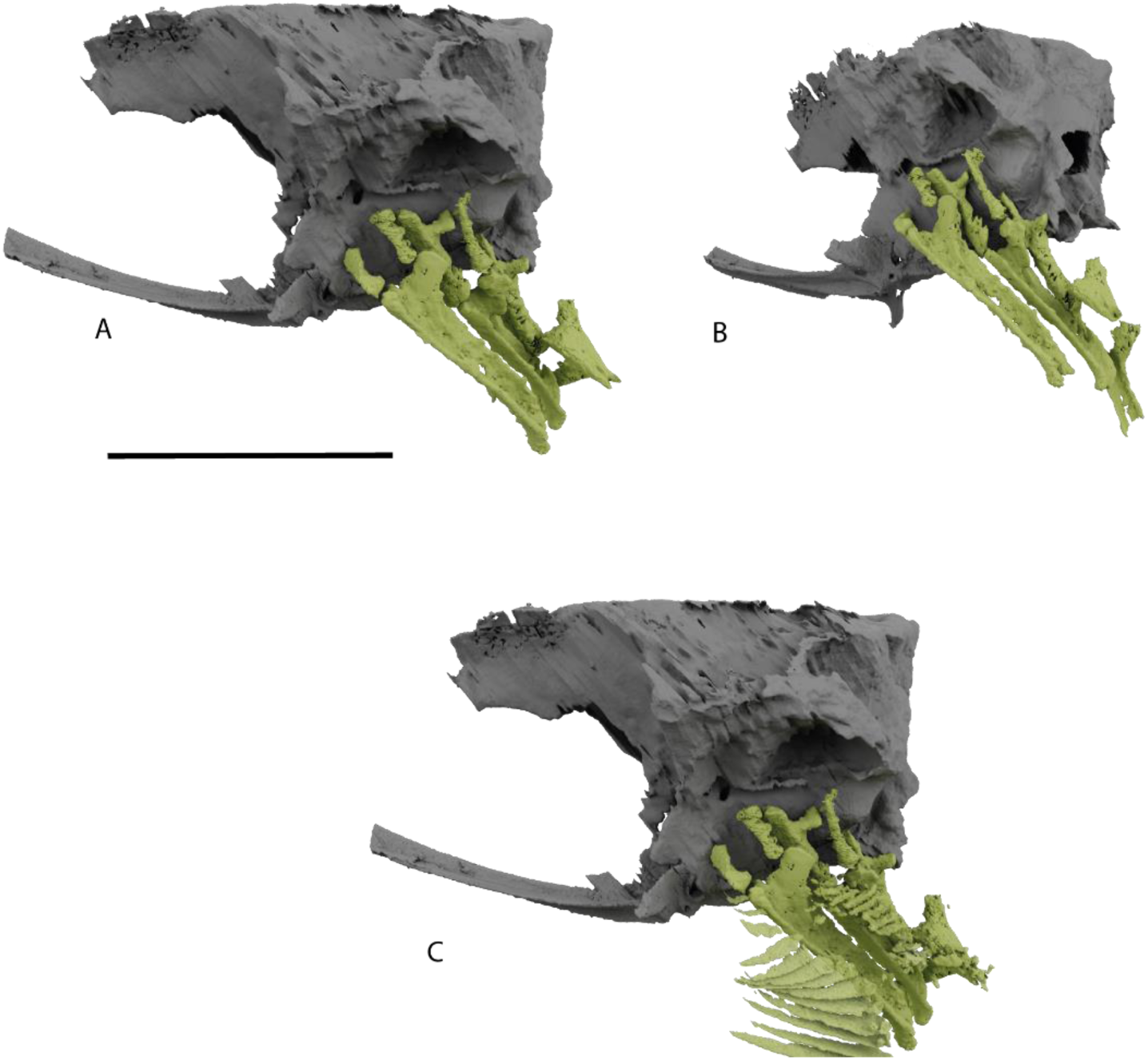
Reconstruction of the gill skeleton of Dorsetichthys bechei NHMUK PV P 1052. Render in (A) right lateral, (B) ventral posterolateral view. (C) Render in right lateral view, with gill rakers and filaments. Scale bar = 10 mm.

**Figure 16:**
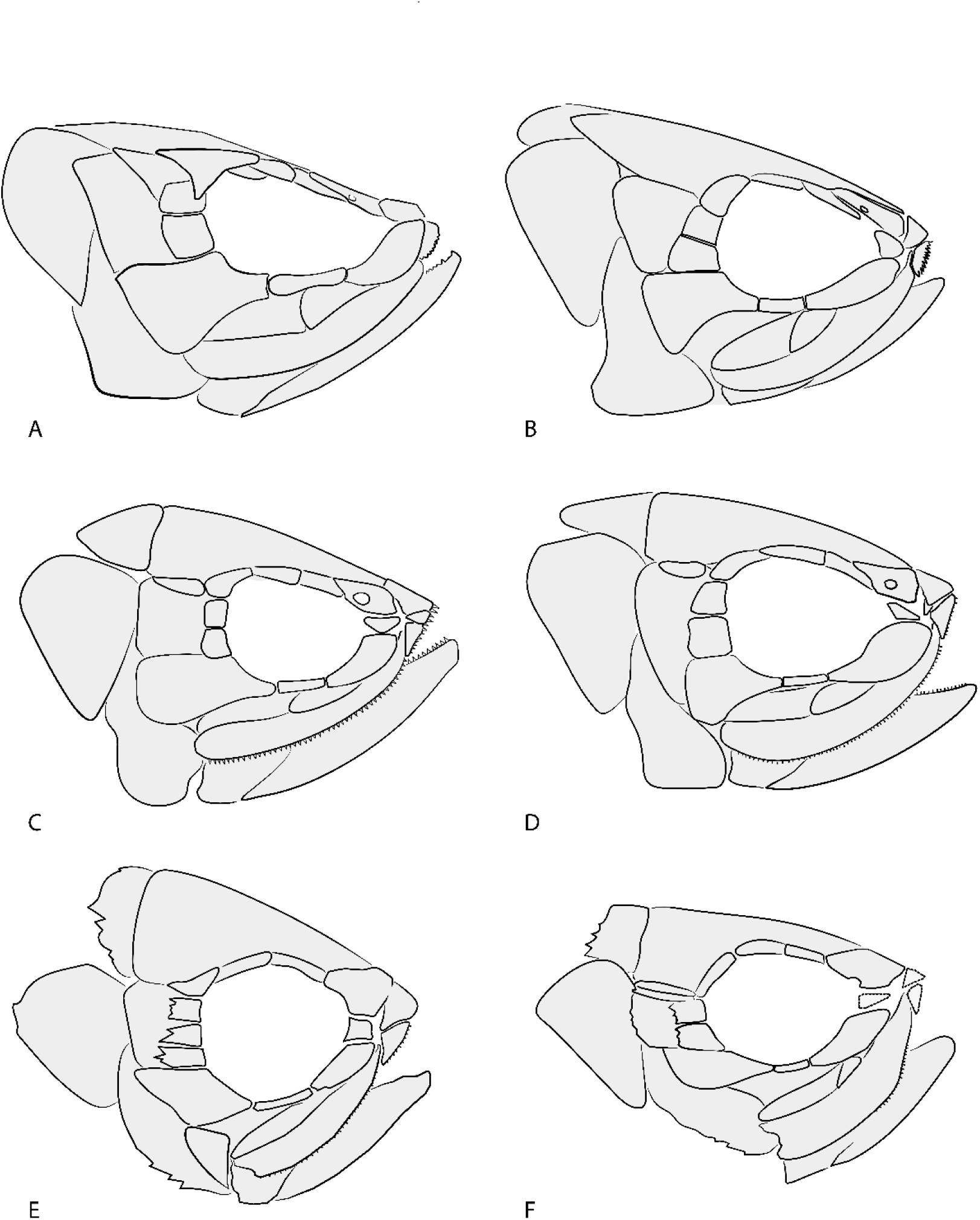
Comparison of dermal skeleton among *Dorsetichthys bechei* and closely related taxa. (A) *Dorsetichthys bechei,* reconstructed based on NHMUK PV P 73995 and NHMUK PV P 1052; (B) *Dorsetichthys bechei,* as previously reconstructed by Nybelin (1966, Fig. 1); (C) *Pholidorhynchodon malzannii* (after Arratia, 2013, Fig. 98); (D) *Parapholidophorus nybelini,* (after Arratia, 2013, Fig. 98); (E) *Pholidoctenus serianus* (after Arratia, 2013, Fig. 7) and (F) *Pholidoctenus sanpellegrinensis* (after Arratia, 2013, Fig. 7). Not to scale.

## Discussion

### Taxonomic coherence of *Dorsetichthys bechei*

Although a key taxon in many investigations of early teleost and neopterygian relationships, the cranial skeleton of *Dorsetichthys* is typically described from an amalgam of three-dimensional but isolated braincase material (Rayner 1948; Patterson, 1975) and abundant but laterally compressed dermal skeletons (Rayner 1948; Nybelin, 1966). Based on comparison to specimens conforming to these disparate modes of preservation, we confirm the identity of NHMUK PV P 73995 as a new specimen of *Dorsetichthys bechei*. The association of external and internal dermal and endoskeletal elements in a single, articulated specimen provides some confidence that past work based on a composite of specimens yielded an accurate picture of anatomy in *Dorsetichthys*. However, there are some anatomical inconsistencies between the material described here and that in past descriptions. For example, the anterior ceratohyal is ungrooved and the posterior ceratohyal does not host branchiostegals, contrasting with conditions in other specimens described by Rayner (1948) and Nybelin (1966). This hints at possible morphological and taxonomic diversity within the material currently referred to *Dorsetichthys bechei*. Future work should target complete specimens that preserve both endoskeletal and dermal elements, and may prove amenable to CT scanning, to investigate further evidence of variation.

### New anatomical insights

CT scanning permits a comprehensive description of the internal and external anatomy of *Dorsetichthys bechei*, allowing us to clarify the specific diagnosis, revise previous descriptions, and provide new insights into its ecology. These updates are spread across the cranium but are particularly concentrated in the upper jaw and palate, lower jaw, and hyoid and branchial arches, and we review these below.

### Upper jaw and palate

In previous descriptions of *D. bechei*, the premaxillae are either described as lacking an anterior connection to the braincase (Rayner, 1948, pg. 40), or the presence or absence of this feature is not remarked upon (Nybelin, 1966; Patterson, 1975). Similarly, descriptions of the ethmoid region (Rayner. 1948; Nybelin, 1966; Patterson, 1975) refer only to partially exposed lateral dermethmoids in two-dimensionally preserved specimens in which internal anatomy cannot be observed or rely on reference to better-preserved ethmoid regions in pholidophorids or leptolepids. Arratia recovers a mobile premaxilla as an unambiguous synapomorphy uniting *Dorsetichthys bechei* with more advanced teleosts (Arratia, 1999, pg. 289; Arratia, 2000, pg. 163), although this appears to be based on an inferred presence in *D. bechei*.

CT scanning of NHMUK PV P 73995 reveals an *in-situ* ethmoid and upper jaw. Both the maxilla and premaxilla have narrow anterior processes that articulate with the ethmoid region to allow protrusion of the dermal elements and mobility of the upper jaw complex. Although three-dimensional braincase material is lacking in pholidophorids *sensu* Arratia (2013), premaxillae appear to be immobile. A fenestra in the ethmoid of “*Pholidophorus” germanicus* (Patterson, 1975, pg. 477), recovered as a possible ankylophorid by Arratia (2017), resembles that seen in *D. bechei* and most likely also permitted movement of the entire upper jaw.

Previous descriptions of the quadrate of *Dorsetichthys bechei* (e.g., Rayner, 1948, Fig. 27A) do not figure or describe a posteroventral process (sometimes referred to as the quadratojugal or quadratojugal process: see Arratia & Schultze, 1991; Taverne, 2018), although it is coded as present in Arratia’s analysis (2013, ch. 78; see discussion in Taverne, 2018) and later included as a diagnostic feature of Dorsetichthyiformes (Arratia, 2017, pg. 22). Here, we confirm the presence of this process in *D. bechei*.

### Lower jaw

The presence of a surangular was included as a diagnostic character of Dorsetichthyiformes by Arratia (2017, pg. 22), despite widespread possession of a surangular in many neopterygians. Curiously, a surangular has not previously been explicitly described as present in *Dorsetichthys bechei*: Rayner (1948: pp. 40) describes the outer surface of the lower jaw as comprising a dentary and angulo-articular, while Nybelin (1966: pp. 363) reports an angulo-supra-angular that cannot be differentiated from the dentary. We find no evidence for a surangular in *D. bechei* and have emended the diagnosis accordingly.

A well-developed postarticular process of the lower jaw is also cited as present in *D. bechei*, and indeed one of the features uniting it with more derived teleosts (Arratia 2013, 2017). However, the anatomical condition more closely resembles that seen in pholidophorids (e.g. Arratia, 2013, pg. 68, fig. 52; pg. 95, fig. 77C), where it is considered absent. We consider the postarticular process to be absent in *D. bechei*.

Previous evidence concerning the presence or absence of a prearticular in *D. bechei* is conflicting. Rayner (1948) states that it is present, although does not figure or illustrate the lower jaw in medial view, whereas Nybelin (1966) reports its absence. Although the internal surface of the lower jaw does not seem to be explicitly described, Arratia considers the prearticular absent in *D. bechei* and codes it as such in phylogenetic analyses (e.g. Arratia 2013, 2015, 2017); absence of a prearticular is therefore one of the characters supporting a close relationship between *D. bechei* and more crownward teleosts to the exclusion of pholidophorids. Here, we describe a prearticular on the lower jaws of both NHMUK PV P 73995 and NHMUK PV P 1052.

### Hyoid and branchial arches

Only limited aspects of the hyomandibula and ceratohyal have previously been described. We describe these elements plus the interhyal, symplectic, and hypohyal in more detail. In both specimens, the anterior ceratohyal is smooth, and in NHMUK PV P 1052, the posterior ceratohyal does not appear to support branchiostegals. This contrasts with previous descriptions, where grooved ceratohyals bearing branchiostegals have been reported (Rayner 1948, Nybelin, 1966); the significance of this variation is not yet known. A subset of teleosts possess paired dorsal and ventral hypohyals. Although Arratia (1999) mentions that only a single hypohyal is present in *D. bechei*, and that it is unknown whether it is pierced by the hyoidean artery, paired dorsal and ventral hypohyals are later reported as a synapomorphy uniting some subset of “advanced teleosts” and *Dorsetichthys* Arratia (2013, 2017). We find evidence for only a single pair of hypohyals in *D. bechei*, which are pierced by the hyoid artery.

Both dorsal and ventral components of the gill skeleton of *D. bechei* support an array of fine rakers and filaments. Similar structures associated with the hyomandibula provide evidence of a pseudobranch. The presence of reduced teeth on the dentary and maxilla in combination with these elaborate gill rakers suggest that *D. bechei* was capable of suspension feeding facilitated by the mobile upper jaw, a feeding mode first alluded to by Rayner (1948). While this feeding mode in stem teleosts is more commonly associated with the giant suspension feeding pachycormids (Friedman *et al.,* 2010, 2013; Schumacher *et al*., 2016; Gouiric-Cavalli, 2017; Dobson et al., 2019), *Dorsetichthys* may have been analogous to small-bodied suspension feeders such as herring and menhaden. Herring exhibit both particle biting (selective capture of individual particles) and filtering (nonselective, passive feeding on any particles in their path), depending on the concentration of prey (Gibson & Ezzi, 1990). As a limited number of small teeth are present on the anterior portion of the dermal jaws of *D. bechei*, a similar facultative suspension feeding ecology seems likely.

### Circumorbital series

NHMUK PV P 73995 and NHMUK PV P 1052 provide some evidence for intraspecific variation in the circumorbital series of *Dorsetichthys*. A second (accessory) suborbital is present posterior to the dermosphenotic in NHMUK PV P 73995, and both NHMUK PV P 73995 and NHMUK PV P 1052 possess three supraorbital bones, rather than the previously reported two; although a third supraorbital was recognised in NHMUK PV P 1052 by Nybelin, it was interpreted as damage sustained during ontogenetic development (Nybelin, 1966, pg. 360). Similar variations in the circumorbital series have been recorded in multiple genera of pholidophorids: *Pholidorhynchodon malzannii* and *Parapholidophorus nybelini* may possess either two or three supraorbitals; the number of supraorbitals may vary on left and right of the same specimen in *P. nybelini*; and a second (accessory) suborbital is also found in some individuals of *P. malzannii, P. nybelini*, and *P. gervasuttii* (Arratia, 2013).

## Implications for relationships

### Dorsetichthys bechei and Pholidophoridae

The wastebasket status of “pholidophorids” (*sensu* Woodward) has presented a significant barrier both for accurate taxonomy and reconstructing patterns of relationships on the teleost stem (Patterson, 1977; Arratia, 2000). Arratia’s (2013) detailed monographic work established a revised diagnosis and set of phylogenetically derived characters upholding the monophyly of a subset of the assemblage. Pholidophoridae (*sensu* Arratia) is diagnosed by a combination of 17 characters, of which six are autapomorphic: absence of an interfrontal suture; tendency for all skull roofing bones to form a fully fused plate; supraorbital and otic canals with tubules that may open to the surface; preopercle with a notched anteroventral margin; long serrated appendage or interclavicular element covering medial surface of cleithrum; clavicle suturing with anteroventral margin of cleithrum; and elongate, leaf like axillary process. Of these proposed autapomorphies, *D. bechei* possesses cephalic sensory canals with small tubules that typically connect to the surface of the bone (e.g. NHMUK PV P 73995, NHMUK PV P 1052), and a notch on the anteroventral margin of the preoperculum is visible in some specimens of *D. bechei* in which this region is unobscured by other ossifications (e.g. NHMUK PV P 38109, NHMUK PV P 3586). The dermal shoulder girdle is poorly preserved and described in *D. bechei*, and the condition of any clavicle or serrated organ is currently unknown.

Monophyly of Pholidophoridae in a phylogenetic context is upheld by two uniquely derived characters: a skull roof with all bones fused into a single plate, and an extrascapular with a large a rollover bony layer at the anterior margin (Arratia 2013, 2017), features both absent in *D. bechei*. The clade is further supported by a large number of homoplastic characters (Arratia 2013: seven; Arratia 2017: 12), a handful of which are present in *D*. *bechei* and a wider sample of teleosts and neopterygians. These variable characters include an orbital region of the skull roof narrower than the postorbital region (we note that the difference in orbital vs postorbital skull roof width in NHMUK PV P 73995 (Figure 1) resembles that seen in pholidophorids), a bony ridge on the dentary separating the splenial and post-splenial regions, pelvic axillary processes, and diplospondyly in the mid-caudal vertebrae. As we find no evidence for a surangular in *D. bechei*, the condition of the coronoid process being contributed to by both the surangular and dentary is no longer shared by *D. bechei* and pholidophorids.

Overall, we find evidence to uphold the removal of *D. bechei* from pholidophorids, but although suggest the diagnosis of Pholidophoridae be revised to clarify characters that are not autapomorphic.

### *Dorsetichthys bechei* and Pholidophoriformes

Arratia’s (2017) review of pholidophorids finds evidence for *Eurycormus* as the sister taxon to Pholidophoridae, together comprising the order Pholidophoriformes. The monophyly of this order is supported by a single unique character and six homoplastic ones. The condition of the uniquely derived character, a clavicle articulating with the anteroventral margin of cleithrum, is unclear in *D. bechei* (see above), but is coded as inapplicable in the matrix of Arratia (2017) and optimised as absent by the parsimony analysis. For all other characters that can be assessed, *D. bechei* displays the contrasting state to Pholidophoriformes. On this basis, we agree with the exclusion of *D. bechei* from Pholidophoriformes.

### *Dorsetichthys bechei* and crownward teleosts

The position of *D. bechei* relative to more crownward teleost nodes is more difficult to determine, with conflicting evidence to support previous hypotheses. All teleosts crownward of Pholidophoriformes (sensu Arratia 2017) are recovered in a trichotomy by Arratia (2017), which represents the most recent phylogenetic analysis to include a substantial assemblage of crown and stem teleosts inclusive of pholidophoriforms. This trichotomy subtends the node uniting *Ichthyokentema*, Ankylophoridae and *D. bechei* plus all remaining teleosts, with *Ichthyokentema* assuming different positions in different most parsimonious trees (Arratia 2017). In Arratia’s strict consensus tree, two uniquely derived characters support this node: presence of a supraoccipital ossification in the braincase (Arratia 2017: c. 16) and presence of an elongate posterodorsal/posteroventral process of the quadrate (Arratia, 2017, c: 86). *D. bechei* shows both of these anatomical states. Seven additional homoplastic characters are also recovered at this node. The anatomy in *D. bechei* is consistent with three of these states (Arratia 2017: c.87, c.151, c.187), variably consistent with one (Arratia 2017: c.149), conflicts with two (Arratia 2017: c.167, c.171), and the condition of one character is unknown (Arratia 2017: c.123). The updated anatomical description of NHMUK PV P 73995 and NMHUK PV P 1052 does little to change our understanding of these characters in *D. bechei*, other than to confirm the presence of a posterodorsal process of the quadrate (Arratia 2017: c.86), separated from the main body of the bone by a distinct notch (Arratia 2017: c.87).

Several additional characters previously reported by Arratia (2013) as having a bearing on stem teleost interrelationships are not optimised in the strict consensus tree, but can be examined in individual most parsimonious trees (Arratia 2017: fig. 10). Many of these characters are homoplastic, but several are particularly pertinent. Absence of a prearticular in the lower jaw (Arratia 2017: c.76) is recovered as supporting either all teleosts crownward of Pholidophoriformes (Arratia 2017: fig. 10A) or as *Ichthyokentema* plus all remaining teleosts (Arratia 2017: fig. 10B); in the latter tree, it is a uniquely derived character. However, we find evidence for a prearticular in *D. bechei* (contra Arratia). A posttemporal with a distinct process to articulate with the cranium (Arratia 2017: c.119) is also recovered as a uniquely derived character in the latter tree, but appears to be absent in *D. bechei* (contra Arratia). Other characters are variable, although we report that a surangular is absent in *D. bechei*; presence of this character, which was previously considered to be present in *D. bechei*, recovered as supporting the clade comprising *Ichthyokentema* and *D. bechei* plus all remaining teleosts (Arratia 2013, 2017).

The placement of *Dorsetichthys*, as sister taxon to *Leptolepis*, Ascalaboidae, Varasichthyidae and crown teleosts (Arratia, 2017: fig. 9) is supported by five homoplastic characters. Our revised description challenges current understanding of the condition of two of these in *D. bechei*. We find no evidence for a well-developed postarticular process of the lower jaw (coded as present in Arratia 2013: c.64 and 2017: c. 72). We also find evidence for a single hypohyal, rather than the dorsal and ventral hypohyals optimised at this node (coded as inapplicable in Arratia 2013: c.82 and 2017: c.91). Furthermore, two of the remaining homoplastic characters, the posterior region of infraorbital 3 extending below the suborbital bone to reach the anterior margin of the preopercle (Arratia 2017: c.47) and a pelvic axillary process (Arratia 2017: c.133) both occur in pholidophorids.

This uncertainty reflects broader instability in patterns of relationship on the teleost stem, as well as difficulties in recovering a monophyletic teleost total group encompassing both extant and fossil taxa on the basis of skeletal characters (e.g. Bardack, 1965; Lehman,1966; Patterson, 1977; Patterson & Rosen, 1977; de Pinna, 1996; Arratia & Schultze, 1990; Arratia, 2001, 2016; Grande, 2010; López-Arbarello & Sferco 2018). CT-based redescriptions of key taxa are one way to addressing this issue as a way of uncovering internal anatomy, including of the neurocranium, palate, and gill skeleton. These techniques are yet to be applied to many stem teleost taxa, most notably aspidorhynchids, pachycormids (but see Dobson et al. 2021) and pholidophorids. Targeted description of these taxa, with a focus on identifying new, robust characters, should be a priority for future investigations, as well as a revised and expanded phylogeny.

## Conclusions

This paper presents a new, previously undescribed specimen of *Dorsetichthys bechei.* Through high-resolution CT scanning, we present a detailed description of both the external dermal skeleton and internal endoskeleton from the same individual for the first time for this important genus. We have identified several anatomical features contrasting with previous descriptions, such as an anterior ceratohyal lacking a groove for the afferent hyoid artery, absence of a surangular, and additional elements of the circumorbital series. We have also confirmed the presence of undescribed but presumed traits, including a mobile premaxilla and quadratojugal process. The presence of elaborate gill rakers, in combination with reduced dentition of the jaws, also supports a new ecological interpretation for *D. bechei*: one of facultative filter feeding. Our new anatomical data support a non-pholidophorid affinity for *D. bechei*. However, the absence of key features that support more crownward nodes on the teleost stem, such as a well-developed postarticular process of the lower jaw, dorsal and ventral hypohyals, and loss of the prearticular, indicate that the position of *D. bechei* relative to the teleost crown is still uncertain. These data pave the way for future expanded phylogenetic reanalyses interrogating patterns of relationship on the teleost stem.

## Acknowledgements

We thank Emma Bernard and Zerina Johanson (NHMUK) for specimen access and Brett Taylor for performing CT scanning. M. Colfer (University of Oxford) carried out preliminary segmentation and interpretation of the Mimics files. S.G. was supported by a Royal Society Dorothy Hodgkin Research Fellowship (no. DH160098). J.A. was supported by a Royal Society Research Grants for Research Fellows (no. RGF\R1\181021) to S.G. William Sanders (Chief Vertebrate Preparator, University of Michigan Museum of Paleontology) removed matrix from NHMUK PV P 73995 to improve CT results. This study includes data produced in the CTEES facility at University of Michigan, supported by the Department of Earth and Environmental Sciences and College of Literature, Science, and the Arts. We would also like to thank the organising committee of *Progressive Palaeontology 2023* (Amber Wood-Bailey, Matt Dempsey and Samuel Cross) for awarding this project Best Talk and offering this fees-waived publication with PeerJ.

## Data availability

For review, segmentation files and surface files are available via temporary Dropbox links:

NHMUK PV P 73995 .mcs file: https://www.dropbox.com/scl/fi/i96u50iae7wgm9jdahgs6/Dorsetichthys-bechei-NHMUK-PV-P73995-CLEAN.mcs?rlkey=u2w1zltgzaat29sod1t6yitc0&dl=0

NHMUK PV P 73995 .ply files (zipped): https://www.dropbox.com/scl/fi/s0rp4vv8hx1b1osyeflqo/NHMUK-PV-P73995-PLYs.zip?rlkey=yqs5jenn352nwxp5cuigy5aab&dl=0

NHMUK PV P 1052 .mcs file: https://www.dropbox.com/scl/fi/l84hsvrfr3ge63w8auo55/Dorsetichthys-bechei-NHMUK-PV-P1052-CLEAN.mcs?rlkey=fp9b5yb2jj80i9rlsjhv092nx&dl=0

NHMUK PV P 1052 73995 .ply files (zipped): https://www.dropbox.com/scl/fi/611010buh7713ivrcda0r/NHMUK-PV-P.1052-PLYs.zip?rlkey=2sraeflp63a4occiiqeqvr37j&dl=0

These, along with the tomogram TIFF stacks, will be deposited in a stable repository upon acceptance.

